# Dietary control of peripheral adipose storage capacity through membrane lipid remodelling

**DOI:** 10.1101/2024.10.25.620374

**Authors:** Marcus J. Tol, Yuta Shimanaka, Alexander H. Bedard, Jennifer Sapia, Liujuan Cui, Mariana Colaço-Gaspar, Peter Hofer, Alessandra Ferrari, Kevin Qian, John P. Kennelly, Stephen D. Lee, Yajing Gao, Xu Xiao, Jie Gao, Julia J. Mack, Thomas A. Weston, Calvin Pan, Aldons J. Lusis, Kevin J. Williams, Baolong Su, Daniel P. Pike, Alex Reed, Natalia Milosevich, Benjamin F. Cravatt, Makoto Arita, Stephen G. Young, David A. Ford, Rudolf Zechner, Stefano Vanni, Peter Tontonoz

## Abstract

Complex genetic and dietary cues contribute to the development of obesity, but how these are integrated on a molecular level is incompletely understood. Here, we show that PPARγ supports hypertrophic expansion of adipose tissue via transcriptional control of LPCAT3, a membrane-bound O-acyltransferase that enriches diet-derived omega-6 (*n*-6) polyunsaturated fatty acids (PUFAs) in the phospholipidome. In high-fat diet–fed mice, lowering membrane *n*-6 PUFA levels by adipocyte-specific *Lpcat3* knockout (*Lpcat3*^AKO^) or by dietary lipid manipulation leads to dysfunctional triglyceride (TG) storage, ectopic fat deposition and insulin resistance. Aberrant lipolysis of stored TGs in *Lpcat3*^AKO^ adipose tissues instigates a non-canonical adaptive response that engages a futile lipid cycle to increase energy expenditure and limit further body weight gain. Mechanistically, we find that adipocyte LPCAT3 activity promotes TG storage by selectively enriching *n*-6 arachidonoyl-phosphatidylethanolamine at the ER–lipid droplet interface, which in turn favours the budding of large droplets that exhibit greater resistance to ATGL-dependent hydrolysis. Thus, our study highlights the PPARγ–LPCAT3 pathway as a molecular link between dietary *n*-6 PUFA intake, adipose expandability and systemic energy balance.

## Introduction

White adipose tissue (WAT) is a dynamic organ that remodels its size, cellular composition, and function in response to hormonal and nutritional signals. To handle surplus energy, WAT can expand its storage volume by increasing adipocyte cell size (hypertrophy) and number (hyperplasia)^1-3^. Hypertrophic growth is the dominant plasticity mechanism in adult humans, and this response is coordinated by the master adipogenic regulator PPARγ^4,5^. Human mutations causing lipodystrophies converge on the PPARγ pathway, affecting central nodes in adipogenesis or direct target genes with non-redundant roles in TG biosynthesis, ER phospholipid remodelling and/or lipid droplet (LD) biology^6^. A hallmark of lipodystrophic states is an impaired expansion capacity of WAT during times of excess energy intake, leading to ectopic fat deposition, lipotoxicity and insulin resistance (IR). Hence, it has been proposed that a high degree of WAT plasticity may be beneficial in obesity^7-9^. In fact, several population-level genetic analyses have suggested that limited peripheral adipose storage capacity, rather than obesity per se, is the real culprit linking nutrient overload to the development of IR^10,11^.

Partitioning of dietary calories into WAT for storage as TGs is directed by a complex interplay of genetic, molecular and environmental cues. The role of diet itself in determining adipose storage capacity, however, is comparatively understudied. Chronic overnutrition predisposes to obesity, yet despite decades of study there is little consensus over the merits and detriments of specific dietary nutrients^12,13^. A growing body of evidence points to changes in both the quantity and quality of dietary fats as causative factors in the development of excess adiposity^14-17^. However, the exact molecular pathways through which dietary fatty acids (FAs) of varying acyl chain lengths and unsaturation degrees shape adipocyte function remain largely unsettled. This limited understanding is due, in part, to a dearth of appropriate models for studying ‘gene–diet’ interactions. For example, animal models of inherited lipodystrophy exhibit defects in fat storage and energy balance regardless of diet composition^18^. In addition, the FA profiles of rodent high-fat diets (HFDs) do not closely mirror human diets^19^.

A major trend in human diets is a rise in the percent of energy intake (en%) derived from *n*-6 PUFAs, most notably linoleic acid (18:2*n*-6)^20^. The resulting increase in the *n*-6:*n*-3 FA ratio has gained considerable traction for its implications in obesity and cardiometabolic disease^21^. *n*-6 linoleic and *n*-3 linolenic (18:3*n*-3) PUFAs are derived exclusively from the diet and serve as competitive substrates for the elongase–desaturase pathway in the formation of highly unsaturated arachidonic acid (20:4*n*-6), eicosapentaenoic acid (20:5*n*-3) and docosahexaenoic acid (22:6*n*-3). Recent work has highlighted divergent effects of *n*-6 and *n*-3 PUFAs on body fat mass by virtue of their ability to modulate adipogenesis, energy expenditure (EE), lipolysis and inflammation^22-26^. By contrast, major gaps persist in our understanding of the molecular driver(s) coupling dietary PUFAs to adaptive adipose remodelling and expansion in the face of caloric surplus. Of interest, the selective accrual of *n*-6 arachidonoyl phospholipids in WAT membranes is a conserved feature of obesity^27^. However, surprisingly little is known about the molecular circuitry that enables diet-dependent rewiring of the adipocyte membrane lipidome and whether such adaptations are of biological importance for proper TG storage and tissue expansion.

Here we show that PPARγ integrates dietary fat intake and adipose storage capacity via transcriptional control of lysophosphatidylcholine acyltransferase 3 (LPCAT3), an ER-resident enzyme that enriches *n*-6 PUFAs in the membrane lipidome. Genetic and dietary interventions highlight the importance of membrane *n*-6 PUFA levels for achieving optimal adipocyte hypertrophic capacity in the setting of diet-induced obesity (DIO). Our study provides a conceptual framework for understanding the effects of *n*-6 PUFAs on WAT plasticity and may inform dietary and therapeutic anti-diabetic strategies.

## Results

### The PPARγ-LPCAT3 axis shapes the adipocyte membrane lipidome

The membrane–bound O-acyltransferase (MBOAT) family member Lpcat3 enriches *n*-6 PUFAs in phosphatidylcholine (PC) and phosphatidylethanolamine (PE), the most abundant phospholipids^28,29^. We found that Lpcat3 was abundantly expressed in fat and was the only LPCAT isoform responsive to PPARγ agonism in murine inguinal WAT (iWAT; Fig. 1a and Extended Data Fig. 1a-c). In addition, its expression levels rose during the course of 10T1/2 adipocyte differentiation (Extended Data Fig. 1d). ChIP experiments confirmed PPARγ binding to a conserved PPAR response element (PPRE) ∼114 bp proximal to the *Lpcat3* transcriptional start site (Fig. 1b and Extended Data Fig. 1e). CRISPR/Cas9 editing of this putative PPRE (**△**PPRE), but not of the known LXRE^30^, abolished the induction of Lpcat3 during differentiation (Extended Data Fig. 1f-h). This editing did not cause general defects in adipogenesis or PPARγ binding to other target genes (Extended Data Fig. 1f-h).

**Fig. 1.**
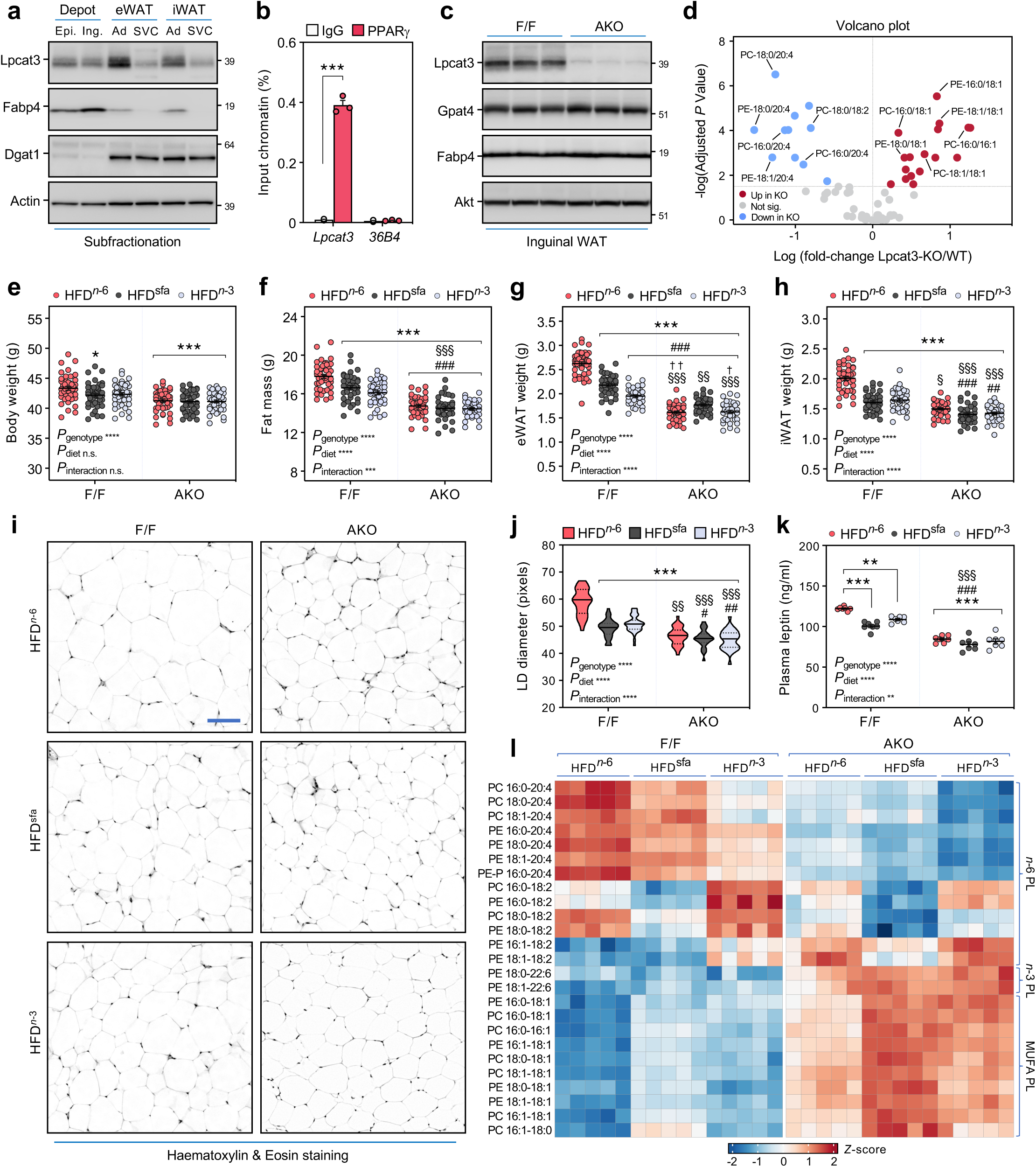
Dietary *n*-6 PUFAs enhance WAT plasticity via the PPARγ-Lpcat3 pathway. (**a**) Immunoblot analysis of LPCAT3 protein levels in whole fat depots, or in adipocyte (Ad) and stromal vascular cell (SVC) membrane fractions of eWAT and iWAT depots. (**b**) ChIP-qPCR analysis of PPARγ binding to the *Lpcat3* promoter in 10T1/2 adipocytes (*n* = 3). (**c**) Immunoblot analysis of LPCAT3 protein levels in iWAT lysates of NCD-fed control and *Lpcat3*^AKO^ mice. (**d**) Volcano plot depicting upregulated (red) and downregulated (blue) PC/PE species in iWAT of 18-week-old NCD-fed *Lpcat3*^AKO^ mice *vs*. controls. (**e**) Body weight and (**f**) EchoMRI analysis of fat mass in 18-week-old control and *Lpcat3*^AKO^ mice fed the indicated 60% HFDs for 10 weeks (*n* = 36–45/group). (**g**,**h**) Wet weights of (**g**) eWAT and (**h**) iWAT from control and *Lpcat3*^AKO^ mice fed the indicated HFDs (*n* = 36–45/group). (**i**) Rescaled H&E stainings of iWAT sections (scale bar, 100 µm) and (**j**) quantification of fat cell diameter (*n* = 6/group). (**k**) Circulating leptin levels in *Lpcat3*^AKO^ mice and controls fed the indicated HFDs for 10 weeks (*n* = 7/group). (**l**) Shotgun-lipidomic analysis and hierarchical clustering of abundant PC/PE species in the inguinal adipocyte fraction from HFD-fed control and *Lpcat3*^AKO^ mice (*n* = 5/group; nomenclature a:b, c:d, where a/c reflect carbon-chain length and b/d the number of unsaturated bonds in *sn*-1 and *sn*-2 aliphatic acyl-chains). Data are presented as mean ± SEM. \*\*\**P* < 0.001 using multiple t-tests with Holm-Sidak’s correction (**b**), adjusted *P*<0.05 using Welch’s *t*-tests with Holm-Sidak’s correction for multiple comparisons on average centered-log ratio (CLR) transformed values for each genotype (**d**), \**P* < 0.05, \*\**P* < 0.01, \*\*\**P* < 0.001 vs. HFD*^n^*^-6^-fed controls; ^#^*P* < 0.05, ^##^*P* < 0.01, ^###^*P* < 0.001 vs. HFD^sfa^; and ^§^*P* < 0.05, ^§§^*P* < 0.01, ^§§§^*P* < 0.001 vs. HFD*^n^*^-3^, same genotype; **^†^***P* < 0.05, **^††^***P* < 0.01 vs. HFD^sfa^-fed *Lpcat3*^AKO^ mice; using two-way RM ANOVA with Tukey’s multiple comparisons test (**e-h**,**j-k**).

To directly address a possible role for LPCAT3 in determining membrane *n*-6 PUFA levels in adipose tissue, we generated a fat-specific *Lpcat3* knockout (AKO) mouse model (Fig. 1c and Extended Data Fig. 2a,b). LPCAT3 loss-of-function was evaluated by shotgun lipidomic analysis of inguinal adipocyte fractions isolated from *Lpcat3*^AKO^ mice and F/F littermate controls. When fed a normal chow-diet (NCD), linoleoyl-PC was abundant in iWAT of control mice, whereas the proportion of arachidonoyl-PC was comparatively minor (Extended Data Fig. 2c). The 18:2*n*-6 and 20:4*n*-6 fatty acyl-chains were more evenly distributed in PE (3:2 ratio, ∼50% of total PE). LPCAT3 deletion caused a prominent deficit of arachidonate (∼60–80%) in PC/PE, but not in adipocyte TGs (Fig. 1d and Extended Data Fig. 2c). We detected reduced linoleoyl-PC (18:0/18:2-PC) in *Lpcat3*^AKO^ iWAT, which was accompanied by an apparent compensatory increase in the abundance of PC/PE subspecies containing palmitoleoyl (16:1*n*-7) and oleoyl (18:1*n*-9) monounsaturated fatty acyl (MUFA) chains (Fig. 1d and Extended Data Fig. 2c). Despite this lipidomic remodelling, no overt histological, functional or gene-expression changes were observed in anatomically distinct *Lpcat3*^AKO^ fat depots (Extended Data Fig. 2c-k).

The loss of LPCAT3 in adipocytes was recently reported to promote systemic insulin sensitivity^31^. We confirmed that iWAT of NCD-fed *Lpcat3*^AKO^ mice had slightly elevated Akt/PKB phosphorylation after an i.p. injection of insulin (Extended Data Fig. 3a). In our model, we found no differences in insulin-stimulated adipocyte glucose disposal, as assessed by PET/CT imaging of ^18^F-FDG uptake (Extended Data Fig. 3b,c). In agreement, glucose and insulin tolerance were comparable between *Lpcat3*^AKO^ mice and their wild-type counterparts (Extended Data Fig. 3d,e). Lastly, metabolic phenotyping showed that the respiratory exchange ratios (RER) were similar between the between the genotypes Extended Data Fig. 3f), indicating no evident changes in carbohydrate utilization.

### The diet-dependent composition of fat cell membranes determines adipose expandability

We next employed a combination of dietary interventions with genetic backgrounds to understand the impact of membrane *n*-6 PUFA levels on adaptive adipose remodelling and expansion. To this end, control and *Lpcat3*^AKO^ mice were fed isocaloric HFDs [60 kcal% fat] with varying proportions of 18:2*n*-6 PUFAs. As our baseline, we opted for the widely used HFD (D12492, Research Diets) containing a relatively high en% from 18:2*n*-6 [HFD*^n^*^-6^; 37:36:27 sfa:mufa:pufa, *n*-6:*n*-3 ratio ∼14:1]. We designed custom two diets in which 18:2*n*-6 PUFA content was partially substituted with either long-chain saturated fats [HFD^sfa^; 59:34:7 sfa:mufa:pufa, *n*-6:*n*-3 ratio 3:1] or 18:3*n*-3 [HFD^*n*-3^; 37:36:27 sfa:mufa:pufa, *n*-6:*n*-3 ratio 1:1]. The dietary FA profiles were confirmed by gas chromatography-mass spectrometry (GC-MS) analyses (Extended Data Fig. 4a).

After 10 weeks of high-fat feeding, the body weights of the control groups trended higher than those of *Lpcat3*^AKO^ mice, most notably on HFD*^n^*^-6^ (Fig. 1e and Extended Data Fig. 4b). This was not attributable to differences in food intake (Extended Data Fig. 4c). EchoMRI analysis revealed a marginal genotype effect on lean body mass regardless of diet (Extended Data Fig. 4d). Of interest, body fat mass gain in controls was strongly influenced by dietary *n*-6 PUFA content, with HFD*^n^*^-6^ provoking greater adiposity than HFD^sfa^ or HFD^*n*-3^ (Fig. 1f). The *Lpcat3*^AKO^ groups exhibited less body fat mass accrual than obese controls and were refractory to changes in dietary lipid composition. The pattern in body fat distribution across the genotypes and dietary regimens aligned with the differences in wet weight of the major WAT depots (Fig. 1g,h), with the greatest degree of expansion observed in HFD*^n^*^-6^-fed controls. Quantification of iWAT cross-sectional areas showed that the increase in fat pad weight was due to adipocyte cell hypertrophy (Fig. 1i,j), and that this was contingent on LPCAT3 expression and *n*-6 PUFA intake. Moreover, the hypertrophic capacity of WAT directly correlated with plasma leptin levels (Fig. 1k).

We next utilized shotgun-lipidomics to interrogate the efficacy of our dietary interventions in lowering *n*-6 PUFA levels in iWAT membranes. Hierarchical clustering uncovered a robust gradient distribution of arachidonoyl-phospholipids across the diets and genotypes (Fig. 1l and Supplementary Table 1). Relative to NCD-fed mice, the greatest 20:4*n*-6 fatty acyl-chain enrichment was observed in HFD^*n*-6^-fed controls. This was particularly evident for the PE lipid class, with arachidonoyl PE (18:0/20:4) emerging as the dominant species. As expected, *n*-6 PUFA abundance in iWAT membranes of HFD^sfa^-fed control mice was limited by reduced dietary provision. By contrast, HFD^*n*-3^-feeding specifically prevented the obesity-related accrual of arachidonoyl-phospholipids in iWAT (Fig. 1l), most likely due to competition of 18:3*n*-3 with 18:2*n*-6 FAs for enzymes of the elongation/desaturation pathway. In support of this notion, linoleoyl phospholipids were enriched in iWAT of HFD^*n*-3^-fed controls (Fig. 1l). In mice lacking Lpcat3, there was a prominent deficit in arachidonoyl phospholipids on all diets. Membrane 18:2*n*-6 acyl-chains were at least partially preserved via their incorporation in atypical PC/PE species (*e.g.,* PC-18:1/18:2, PE-18:1/18:2). The proportion of DHA-phospholipids was unaffected by dietary lipid manipulation, even upon ample *n*-3 PUFA provision (Fig. 1l and Supplementary Table 1). 18:3*n*-3 and 20:5*n*-3 were not consistently detected in the membrane lipidome but were represented abundantly in PUFA-derived oxylipins and in adipocyte TGs (Extended Data Fig. 4e and Supplementary Table 1).

Reduced fat mass can be due to a primary increase in EE or an underlying defect in storage capacity. We observed a marked increase in the size and weight of *Lpcat3*^AKO^ livers relative to HFD^*n*-6^-fed controls (Fig. 2a,b), consistent with a partial lipodystrophic phenotype in which FAs are diverted from adipose tissue to liver^32^. *Lpcat3*^AKO^ livers were steatotic regardless of dietary composition, as evidenced by the build-up of hepatic TGs and cholesterol esters (Fig. 2a-c). A similar trend was detected for plasma TGs and ALT/AST levels (Extended Data Fig. 4f-i). Importantly, control groups fed *n*-6 PUFA-depleted HFDs also displayed enlarged fatty livers (Fig. 2a-c), phenocopying the *Lpcat3*^AKO^ model. This was not due to impaired hepatic VLDL-TG secretion or β-oxidation (Extended Data Fig. 4j,k), indicating that fatty liver development was secondary to limited storage capacity in adipose tissue. Finally, the protection against ectopic fat deposition in HFD^*n*-6^-fed controls tracked with greater insulin tolerance (Fig. 2d,e). These data suggest that *n*-6 PUFAs enhance adipose-tissue plasticity via the PPARγ-LPCAT3 axis.

**Fig. 2.**
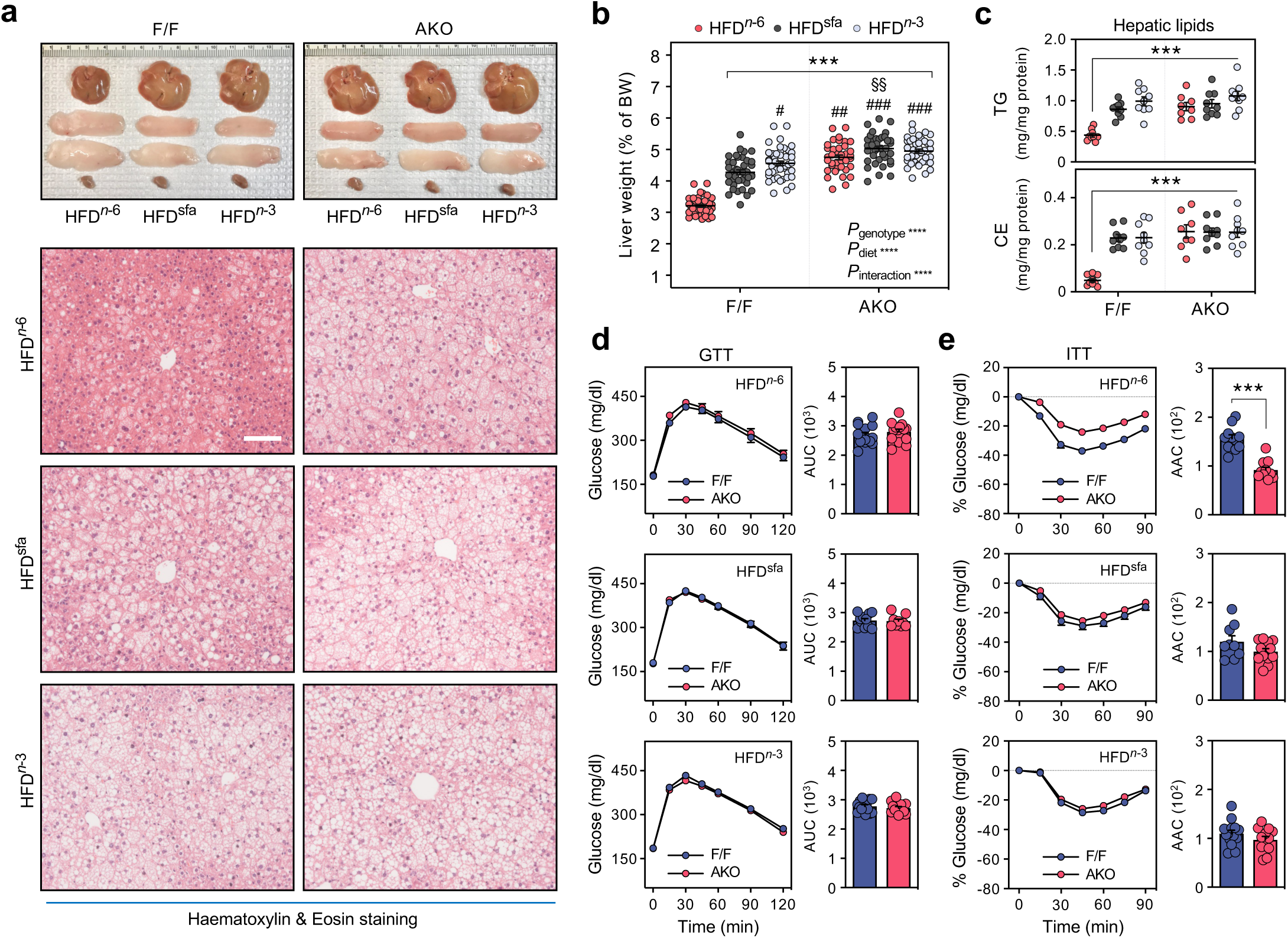
Lowering *n*-6 PUFA levels in WAT membranes predisposes to lipodystrophy. (**a**) H&E staining of liver sections (scale bar, 50 µm) and macroscopic appearance of the liver and distinct fat depots (top to bottom: iWAT, eWAT, and iBAT) from 18-week-old *Lpcat3*^AKO^ mice and controls fed the indicated HFDs for 10 weeks. (**b**) Liver wet weights (% of body weight, BW; *n* = 36–45/group) and (**c**) hepatic TGs and cholesterol esters (CE) in HFD-fed control and *Lpcat3*^AKO^ mice (*n* = 8–9/group). (**d**) Glucose tolerance (1 g kg^-1^) and (**e**) insulin tolerance tests (1.25 U kg^-1^) performed on control and *Lpcat3*^AKO^ mice fed the indicated HFDs for 10-12 weeks (*n* = 9–15/group). Data are presented as mean ± SEM. \*\*\**P* < 0.001 vs. HFD*^n^*^-6^-fed controls; ^#^*P* < 0.05, ^##^*P* < 0.01, ^###^*P* < 0.001 vs. HFD^sfa^; ^§§^*P* < 0.01 vs. HFD*^n^*^-3^, same genotype; by two-way RM ANOVA followed by Tukey’s multiple comparisons test (**b**-**c**) or two-tailed *t*-test analysis of the area under (**d**) or above the curve (**e**).

### An adaptive response in obese *Lpcat3*^AKO^ iWAT offsets unbridled lipolysis via TG/FA cycling

To investigate the underlying cause of dysfunctional fat storage in the *Lpcat3*^AKO^ model, we conducted a series of lipid radiotracer studies to analyse the kinetics of TG synthesis and hydrolysis in the HFD^*n*-6^-fed genotypes. After 5 weeks on HFD^*n*-6^, there were no changes in the lipid uptake and oxidation rates in fat or other peripheral insulin-target tissues (Extended Data Fig. 5a,b). The secretion of glycerol and NEFAs from obese *Lpcat3*^AKO^ iWAT explants was ∼3-5-fold higher under basal and β3-adrenoreceptor-activated lipolytic states following 5 and 10 weeks of high-fat feeding (Fig. 3a and Extended Data Fig. 5c), suggesting that the storage deficit arose from unbridled TG hydrolysis. Unexpectedly, *Lpcat3*^AKO^ mice had *greater* FA disposal into WAT relative to controls after 10 weeks on HFD^*n*-6^, most notably in inguinal fat depots (Extended Data Fig. 5d). *In vivo* pulse-chase analysis confirmed an accelerated ^14^C-oleate turnover rate in obese *Lpcat3*^AKO^ iWAT (Fig. 3b), indicative of adaptive TG/FA cycling. Befittingly, we detected parallel increases in *de novo* lipogenesis (DNL) and FA β-oxidation in iWAT of HFD^*n*-6^-fed *Lpcat3*^AKO^ mice (Fig. 3c,d). We did not find differences in FA uptake and turnover in brown adipose tissue (BAT) or liver (Fig. 3c,d and Extended Data Fig. 5e,f).

**Fig. 3.**
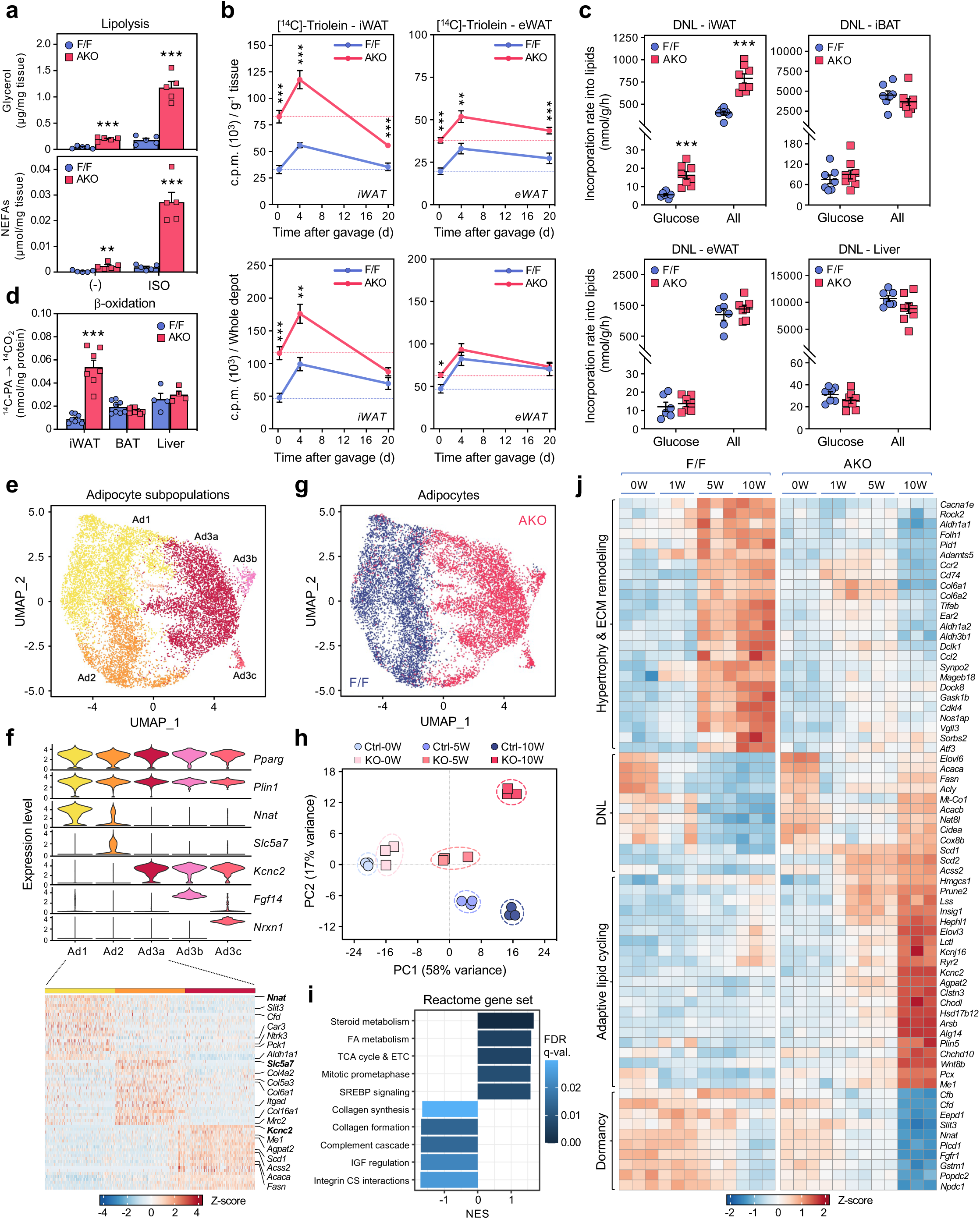
Dysfunctional fat storage and adaptive TG/FA cycling in the obese *Lpcat3*^AKO^ model. (**a**) *Ex vivo* lipolysis in freshly dissected obese iWAT explants from 18-week-old *Lpcat3*^AKO^ mice and F/F controls fed a HFD*^n^*^-6^ for 10 weeks. Released glycerol and NEFAs were analysed under basal and stimulated (2 *µ*M isoproterenol, ISO) lipolytic states (*n* = 5/group). (**b**) ^14^C-oleate uptake and turnover in iWAT and eWAT of *Lpcat3*^AKO^ mice and controls fed a HFD*^n^*^-6^ for 10 weeks (*n* = 6–9/group). Radioactivity (counts per min, CPM) was calculated in extracted lipids from whole-organs 4 h, 4 d, and 20 d post-gavage with ^14^C-Triolein and normalized for g/tissue or whole depot. (**c**) *In vivo* DNL from glucose or all substrates in fat depots and the liver of *Lpcat3*^AKO^ and controls mice fed a HFD*^n^*^-6^ for 10 weeks (*n* = 6–8/group). (**d**) *Ex vivo* β-oxidation rates in crude iWAT, iBAT, or liver lysates, assessed by the conversion of ^14^C-palmitic acid (PA) into ^14^CO_2_ (*n* = 4–7/group). (**e**) UMAP projection of clusters formed by 11,474 adipocytes: sNuc-seq was performed on iWAT depots of control and *Lpcat3*^AKO^ mice fed a HFD*^n^*^-6^ for 10 weeks (*n* = 3, 3). (**f**) Violin plot (clusters as columns, genes as rows) and heatmap of Ad1-3 subtypes. (**g**) UMAP projection of the iWAT transcriptome separated for genotype. (**h**) Principal component analysis (PCA) of iWAT transcriptome in *Lpcat3*^AKO^ mice and controls during the course of high-fat feeding. RNA-seq was performed on age-matched mice that were kept on NCD or subjected to a HFD*^n^*^-6^ for 1, 5, or 10 weeks (*n* = 3/group). (**i**) Gene-set enrichment analysis (GSEA) of up- and downregulated pathways in iWAT of *Lpcat3*^AKO^ mice *vs.* controls fed a HFD*^n^*^-6^ for 10 weeks (normalized enrichment score, NES). (**j**) Heatmap transformation reflecting temporal dynamics of the iWAT transcriptome. Data are presented as mean ± SEM. \**P* < 0.05, \*\**P* < 0.01, \*\*\**P* < 0.001 by multiple *t*-tests with Holm-Sidak’s correction (**a**-**e**).

To understand the molecular underpinnings of altered lipid handling in *Lpcat3*^AKO^ mice, we conducted single-nuclei RNA-seq (sNuc-seq) analyses of iWAT. Adipocytes comprised the largest population of nuclei (Extended Data Fig. 6a,b). Re-clustering broadly separated these nuclei into 3 subpopulations, all of which expressed higher levels of classical pan-adipogenic markers (*e.g.*, *Plin1*, *Lep*, *Cidec*) relative to the stromal vascular cell fraction (Fig. 3e,f and Extended Data Fig. 6c). The Ad1 subtype was defined by the dormancy marker *Nnat*^33^, and was strongly enriched for transcripts encoding lipid storage and growth factor signalling components (*e.g*., *Pck1*, *Ntrk3*, *Cfd*, *Car3*). The Ad2 subtype was characterised by *Slc5a7* and expressed higher levels of genes related to extracellular matrix (ECM) remodelling (*e.g*., *Col5a3*, *Col6a1/2*, *Mrc2*). The Ad3 subtype was defined by the K*^+^* voltage-gated channel *Kcnc2* and was enriched in lipid biosynthetic genes (*e.g.*, *Acss2*, *Agpat2*, *Scd1*). Two smaller Ad3 (Ad3b/c) subtypes separated based on *Fgf14* and *Nrxn1* transcripts, respectively. Genotype clustering revealed that control adipocytes were primarily comprised of Ad1/2 subtypes (Fig. 3g and Extended Data Fig. 6d), befitting active hypertrophic expansion^34^. Remarkably, Lpcat3-deficient adipocytes uniformly clustered in the lipogenic Ad3 subtype (Fig. 3g). Intact DNL enzyme expression in obese *Lpcat3*^AKO^ iWAT was confirmed by immunoblotting analysis (Extended Data Fig. 6e). This was of interest, given that DNL activity in fat correlates with systemic insulin sensitivity and is known to be suppressed in obesity^35-38^.

To expand on our sNuc-seq analyses, we mapped the temporal dynamics of the iWAT transcriptome in response to DIO. The gene-expression profiles of both genotypes progressively diverged over the course of high-fat feeding (Fig. 3h-j). In line with earlier reports^39-41^, iWAT of HFD^*n*-6^-fed controls adopted a classic “hypertrophic” signature comprised of ECM remodelling and TGFβ-related gene programs (Extended Data Fig. 7a and Supplementary Table 2). On the other hand, high-fat feeding potently suppressed lipogenic gene expression in controls, whereas transcript levels of DNL genes in *Lpcat3*^AKO^ iWAT were unaffected by DIO (Fig. 3i,j). We also identified a distinct gene program engaged only in obese *Lpcat3*^AKO^ iWAT, coinciding with the onset of adaptive lipid cycling. Genes in this group were enriched for lipid biosynthetic processes, fatty acid catabolism and cation transport (Extended Data Fig. 7b and Supplementary Table 2). By contrast, transcripts related to creatine and calcium-dependent substrate cycles, which previous work has implicated in EE^42,43^, were unaltered in *Lpcat3*^AKO^ iWAT (Extended Data Fig. 7c). We hypothesized that DIO-induced remodelling in the *Lpcat3*^AKO^ iWAT transcriptome posed a metabolic adaptive response to defective storage capacity. In agreement, key markers of this program were highly upregulated in anatomically distinct WAT depots, but not in BAT (Extended Data Fig. 8a-c). Combined RNA-seq and gene-set enrichment analyses (GSEA) further revealed that dietary *n*-6 PUFA restriction or the loss of CIDEC (a gene linked to human partial lipodystrophy^44^) elicited WAT molecular signatures nearly identical to that in obese *Lpcat3*^AKO^ mice (Extended Data Fig. 9a-c), pointing to the existence of a more broadly conserved adaptation to dysfunctional fat storage.

Animal models of lipodystrophy often manifest “lean and healthy” phenotypes in the face of chronic overnutrition, which has been ascribed to a higher metabolic rate^45^. Meta-analysis of published data from the <u>Met</u>abolic <u>S</u>yndrome in <u>M</u>en study (METSIM; *n* =10,000) revealed negative correlations of key *Lpcat3*^AKO^ markers (including *CHODL*, *PLIN5*, *ACSS2* and *WNT8B*) with BMI, adiposity, plasma TG and IR indices (Extended Data Fig. 9d). When challenged with a prolonged HFD^*n*-6^ regimen, *Lpcat3*^AKO^ mice developed a marked resistance to DIO relative to controls (Fig. 4a,b), coinciding with the onset of the futile lipid cycling program. Indirect calorimetry validated that the plateau in *Lpcat3*^AKO^ body weight gain was due to increased EE, with no changes in locomotor activity (Fig. 4c,d). Of note, this adaptive mode of EE was independent of cell-intrinsic effects of Lpcat3 deletion in brown/beige fat depots (*Lpcat3*^BKO^; Fig. 4e-h). Moreover, HFD^*n*-6^- fed *Lpcat3*^AKO^ mice still displayed a DIO-resistant phenotype when housed at thermoneutrality (Fig. 4i,j) or when crossed onto a *Ucp1^-/–^* (*U1/L3*^DKO^) background (Fig. 4k,l), indicating that the increased metabolic rate was independent of classic WAT “browning” pathways^46^. Accordingly, HFD^*n*-6^-fed *Lpcat3*^AKO^ mice on both *Ucp1^+/+^*and *Ucp1^-/–^* backgrounds maintained a higher core body temperature relative to littermate controls during acute cold challenge (Fig. 4m-o). Our data suggest that obese *Lpcat3*^AKO^ mice engage in an adaptive program of UCP1-independent futile TG/FA cycling in iWAT as a defence against further body weight gain and ectopic fat deposition.

**Fig. 4.**
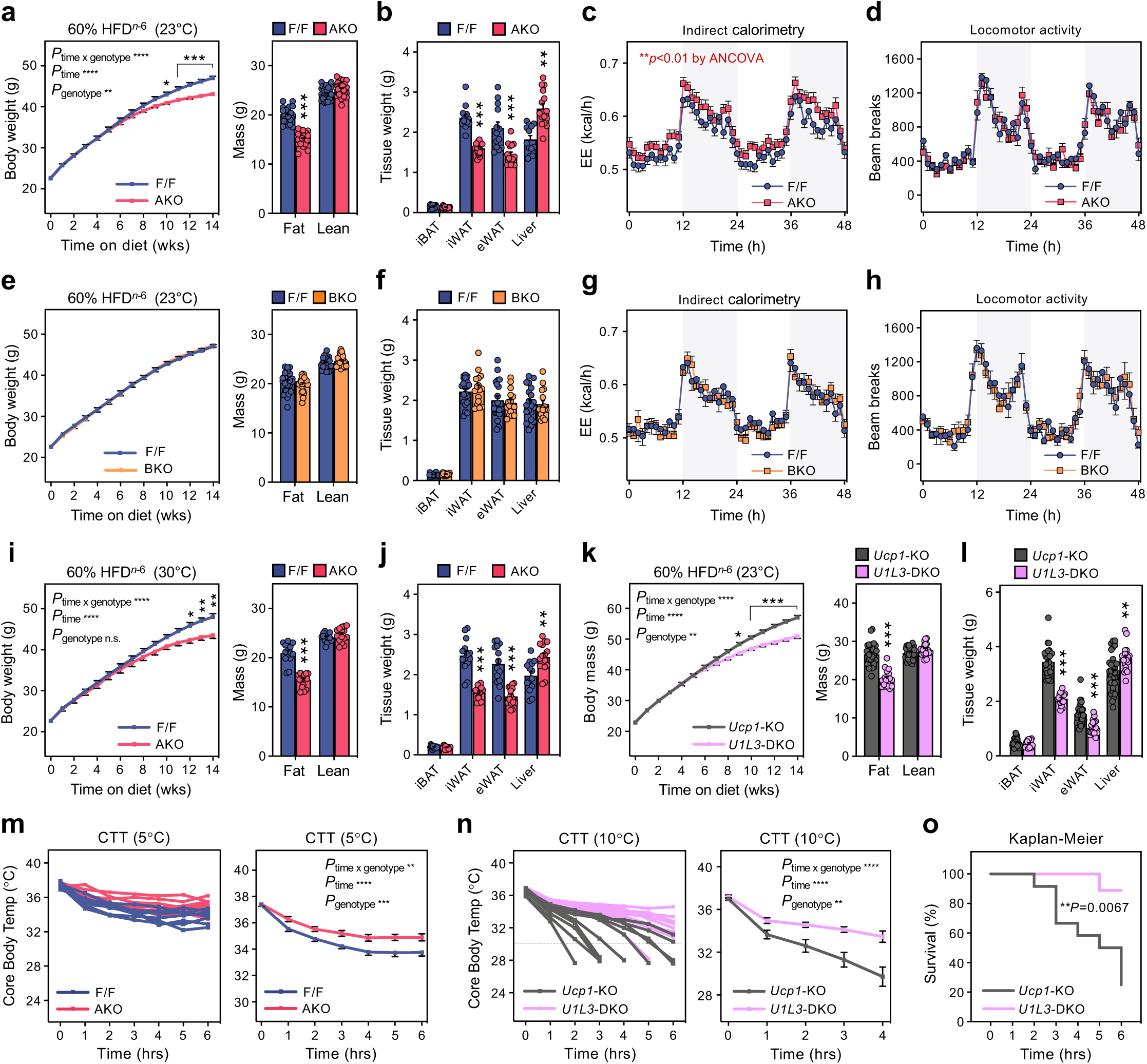
Adaptive futile lipid cycling in iWAT protects *Lpcat3*^AKO^ mice against DIO. (**a**) Body weight (BW) curves and composition of *Lpcat3*^AKO^ mice and controls fed a HFD*^n^*^-6^ for 14 weeks (*n* = 34, 41). (**b**) Wet weight of the indicated tissues in HFD*^n^*^-6^-fed *Lpcat3*^AKO^ mice and controls (*n* = 13, 15). (**c**,**d**) Indirect calorimetry in *Lpcat3*^AKO^ mice and controls fed a HFD*^n^*^-6^ for 12 weeks. (**c**) EE (kcal/h) and (**d**) locomotor activity were monitored over a period of 48 h in Oxymax-CLAMS metabolic cages (12 h light/dark cycles; *n* = 25, 27). (**e**) BW curves and composition of *Lpcat3*^BKO^ mice and controls fed a HFD*^n^*^-6^ for 14 weeks (*n* = 16, 20). (**f**) Wet weight of the indicated tissues in HFD*^n^*^-6^-fed *Lpcat3*^BKO^ mice and controls (*n* = 16, 20). (**g**, **h**) Indirect calorimetry in *Lpcat3*^BKO^ mice and controls fed a HFD*^n^*^-6^ for 12 weeks: (**g**) EE (kcal/h) and (**h**) locomotor activity (*n* = 13, 15). (**i**) BW curves and composition of *Lpcat3*^AKO^ mice and controls housed at thermoneutrality (TN; 30°C) and fed a HFD*^n^*^-6^ for 14 weeks (*n* = 14, 15). (**j**) Wet weight of the indicated tissues in HFD*^n^*^-6^-fed *Lpcat3*^AKO^ mice and controls (*n* = 14, 15). (**k**) BW curves and composition of *Ucp1*^KO^ and *U1L3*^DKO^ mice fed a HFD*^n^*^-6^ for 14 weeks (*n* = 32, 19). (**l)** Wet weight of the indicated tissues in HFD^*n*-6^-fed *Ucp1*^KO^ and *U1L3*^DKO^ mice (*n* = 32, 19). (**m-o)** Cold tolerance tests (CTT) performed on (**m**) control and *Lpcat3*^AKO^ mice (*n* = 6, 6) or (**n**) *Ucp1*^KO^ and *U1L3*^DKO^ mice (*n* = 9, 10) fed HFD*^n^*^-6^ for 10 weeks. Core body temperatures ≤28°C were not included in CTT plots and INSTEAD scored as events for (**o**) Kaplan-Meier survival curve. Data are presented as mean ± SEM. \**P* < 0.05, \*\**P* < 0.01, \*\*\**P* < 0.001 by two-way RM ANOVA with Holm-Sidak’s correction (**a**,**i**,**k**), a two-sided Welch’s *t*-test (**a**-**b**,**i**-**l**), ANCOVA using lean mass as a covariate (**c**), or Gehan-Breslow-Wilcoxon test (**o**).

### Adipocyte LPCAT3 activity optimizes TG storage by converging on LD budding size

To explore the molecular basis of lipolytic dysregulation in the *Lpcat3*^AKO^ model, we generated *Lpcat3*-KO 10T1/2 clones (*L3*_E1/E3; Extended Data Fig. 10a) and stably expressed either GFP control, Lpcat3^WT^, or a catalytically inactive Lpcat3^H374A^ in these lines. As expected, arachidonoyl-phospholipids were readily detected in Lpcat3^WT^-reconstituted adipocytes, with the PE lipid class having a much greater concentration of arachidonate relative to PC (Extended Data Fig. 10b and Supplementary Table 3). Lpcat3^H374A^ and GFP-expressing cells were biased towards incorporating MUFA chains. This lipidomic rewiring did not affect fat cell development *per se*, as the induction of pan-adipogenic markers was comparable between Lpcat3^WT^ and Lpcat3^H374A^-expressing cells (Extended Data Fig. 10c). Strikingly, however, Lpcat3^WT^ re-expression greatly increased LD size and TG storage (Fig. 5a,b). This was dependent on its catalytic activity, since Lpcat3^H374A^ was unable to “rescue” the storage deficit. In agreement, the cell-active Lpcat3 inhibitor (*R*)-HTS-3 blunted TG accrual in Lpcat3^WT^-reconstituted adipocytes and in a parallel wild-type 10T1/2 clone (Fig. 5c, Extended Data Fig. 10d-f, and Supplementary Table 4).

**Fig. 5.**
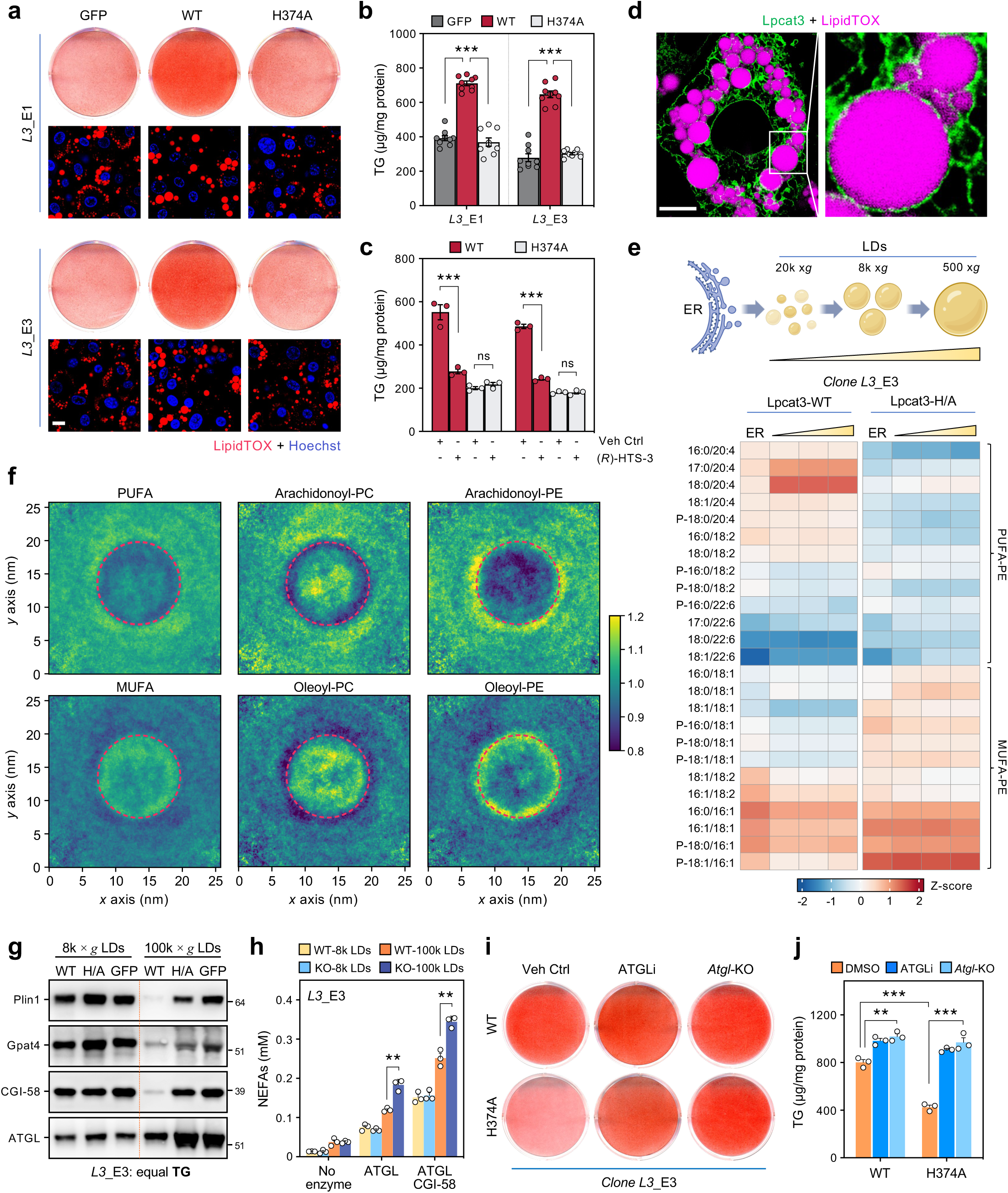
Adipose Lpcat3 activity promotes TG storage by converging on LD budding size. (**a**) Oil red-O (ORO) stainings (top), LipidTOX stainings (bottom: scale bar, 10 µm), and (**b**) TG content in *Lpcat3*-KO 10T1/2 adipocytes stably expressing GFP ctrl, Flag-Lpcat3^WT^, or Flag-Lpcat3^H374A^ (day 7 of differentiation) (*n* = 9/group). *Lpcat3*-null clonal cell lines were made by CRISPR/Cas9 gene editing of exon 1 or 3 (E1/E3). (**c**) TG content in *Lpcat3*-KO adipocytes stably expressing Lpcat3^WT^ or Lpcat3^H374A^ and differentiated in the presence of 0.1% DMSO (vehicle, veh ctrl) or Lpcat3 inhibitor (*R*)-HTS-3 (10 *µ*M) for 5 days (*n* = 3/group). (**d**) Analysis of intracellular Lpcat3-GFP^KKRE^ distribution relative to LipidTOX-stained LDs in *Lpcat3*-KO adipocytes by live-cell confocal imaging (scale bar, 5 µm). (**e**) Shotgun-lipidomic analysis of PE lipid class composition in purified ER membranes and buoyant LD-ER-enriched fractions comprised of small/nascent (20,000 × *g*), medium (8,000 × *g*) or large (500 × *g*) LDs from *Lpcat3*-null adipocytes expressing Lpcat3^WT^ or Lpcat3^H374A^. Data are presented as the distribution of PE molecular species (% of total PE) within each sample. (**f**) 2D lateral density enrichment-depletion maps of MUFA- and PUFA-phospholipid partitioning in a model bilayer that mimics LD budding. CG-MD simulations were carried out on a “blister-shaped” model bilayer/monolayer of 25%SOPC-25%SOPE (MUFA) and 25%SAPC-25%SAPE (PUFA) with a triolein oil core (see Extended Data Fig. 11i). Red circles denote the budding lens. (**g**) Immunoblot analysis of medium (8,000 × *g*) or small/nascent (100k × *g*) LD fractions isolated from *Lpcat3*-null adipocytes stably expressing Lpcat3^WT^ or Lpcat3^H374A^ (day 6 of differentiation). Equal amounts of TG from each condition were loaded onto gel. (**h**) LD-associated lipolytic activity. Medium (8,000 × *g*) and small (100k × *g*) LD fractions were isolated from *Lpcat3*-null adipocytes expressing Lpcat3^WT^ or Lpcat3^H374A^ (day 6 of differentiation). *In vitro* NEFA release was measured 1 h post-incubation of LDs with purified recombinant ATGL ± CGI-58 (*n* = 3/group). (**i**) ORO staining and (**j**) TG content in *Lpcat3*^KO^ or *Lpcat3*/*Atgl*^DKO^ adipocytes stably expressing Lpcat3^WT^ or Lpcat3^H374A^ and differentiated in the presence of 0.1% DMSO (veh ctrl) or Atglistatin (ATGLi, 40 *µ*M) for 7 days (*n* = 3/group). Data are presented as mean ± SEM. \*\**P* < 0.01, \*\*\**P* < 0.001 by multiple *t*-tests with Holm-Sidak’s correction (**b**-**c**,**h**-**j**).

LD formation is initiated when neutral lipids are deposited between the leaflets of the ER bilayer. At a critical concentration, neutral lipids coalesce, forming an oil lens in a process of demixing. The lens grows by amassing TGs and ultimately gives rise to a spherical LD that is enveloped by a phospholipid monolayer derived from the cytosolic face of the ER. Ample evidence suggests that the composition of ER membranes determines LD budding and growth^47-49^. To understand how LPCAT3 affects LD size/stability, we studied its subcellular localization by live-cell imaging. We engineered an Lpcat3 vector harbouring a GFP tag prior its C-terminal ER retention motif. This fusion protein retained full catalytic activity and rescued TG storage when stably expressed in *Lpcat3*-null adipocytes (Extended Data Fig. 10g-i). The LPCAT3 signal displayed a clear reticular pattern that was enriched at the LD periphery (Fig. 5d). Subcellular fractionation confirmed that LPCAT3 was ER-resident and did not redistribute to the LD surface (Extended Data Fig. 11a), in line with its multi-pass membrane-spanning topology^50^. Based hereon, we posited that LPCAT3 activity rewires the fatty acyl-chain profile of local ER regions to facilitate nascent LD assembly. Comparative lipidomics of bulk ER and LD/ER-enriched buoyant fractions demonstrated a high degree of similarity in PC composition (Extended Data Fig. 11b). On the other hand, we observed a notable shift in the distribution of PE molecular species, with LD-associated membranes having a much greater proportion of arachidonoyl-PE relative to ER (Fig. 5e and Supplementary Table 5). This increase was contingent on the expression of LPCAT3 and came at the expense of MUFA-PE.

Prior work has demonstrated that the unique topology of the LD surface is important for cytosol-to-LD protein targeting^51-53^. We therefore performed atomistic molecular dynamics (MD) simulations to compute the impact of LPCAT3 deficiency on the surface properties of model membranes. Increasing the MUFA-to-PUFA ratio in bilayers and ‘LD-like’ trilayer systems did not notably affect the appearance of lipid-packing defects (LPDs; Extended Data Fig. 11c,d). Accordingly, *Lpcat3*-null adipocytes did not exhibit differences in membrane packing, canonical lipolytic receptor signalling pathways or LD protein targeting (Extended Data Fig. 11e-h). We next employed course-grain MD simulations to study the partitioning of arachidonoyl-phospholipids in a mixed bilayer/monolayer system designed to mimic nascent LDs^54^. This model consists of a ‘blister-shaped’ model bilayer that unzips, forming two monolayers to accommodate a triolein oil core (Extended Data. Fig. 11i). 2D lateral density enrichment-depletion maps revealed that arachidonoyl-PC had no preference for the budding lens or the bilayer (Fig. 5f). By contrast, oleoyl-PC uniformly redistributed to the surface monolayers covering the oil lens. Befitting the negative curvature of conically shaped lipids, PE generally partitioned to the interface between the ER bilayer and LD monolayers (Fig. 5f). Like oleoyl-PC, oleoyl-PE was in direct apposition to the triolein oil core, likely owing to their structurally compatible fatty acyl-chain profiles. In support of this notion, arachidonoyl-PE partitioned more towards the outer periphery of the ER–LD interface.

Our combined data suggested that the PE acyl-chain profile at ER–LD junctions is a key determinant of LD size regulation. Electron microscopy of *Lpcat3*-null adipocytes revealed irregular clusters of very small droplets (Extended Data. Fig. 11j), consistent with a defect in budding size. In agreement, small/nascent LD fractions isolated from GFP or Lpcat3^H374A^-expressing cells had higher levels of LD-associated proteins per unit of triglyceride relative to those from Lpcat3^WT^-reconstituted adipocytes (Fig. 5g). This was particularly evident for adipose TG lipase (ATGL) and its co-activator CGI-58, reflecting increased accessibility of the lipolytic machinery to an expanded LD surface area. Accordingly, small/nascent LD fractions isolated from *Lpcat3*-KO adipocytes were more prone to TG hydrolysis following exposure to affinity-purified lipolytic enzymes (Fig. 5g,h). This was contingent on the defect in budding size, as mature droplets had similar LD protein targeting and lipolytic activity (Fig. 5g,h). Finally, suppressing ATGL activity through either genetic or pharmacologic interventions largely rescued TG storage in *Lpcat3*-null adipocytes (Fig. 5i,j and Extended Data Fig. 11k). Collectively, our data indicate that LPCAT3 promotes TG storage by enriching arachidonoyl PE at the ER–LD interface, which favours the budding of large droplets that are less susceptible to cytosolic lipase activity.

## Discussion

The Lands cycle of phospholipid deacylation–reacylation was first described more than 60 years ago^55^. Our study positions Lands’ cycle front and centre in dietary control of peripheral adipose storage capacity. We identify the PPARγ–LPCAT3 axis as an essential link between *n*-6 PUFA intake, adipose expandability, and systemic energy balance. LPCAT3 localizes to ER–LD contacts in adipocytes, where it controls LD size through a mechanism dependent on its catalytic activity. Combined lipidomic and *in silico* analyses further showed that this involves dynamic remodelling of PE fatty acyl-chain composition at the ER–LD interface, which favours the budding of larger droplets that are less prone to ATGL-mediated hydrolysis. Accordingly, mice lacking LPCAT3 manifest defects in adipocyte hypertrophy and TG storage due to unbridled lipolysis, resulting in a lipodystrophic phenotype. Our study delineates a molecular basis for dietary effects on body fat distribution and offers an adipocentric view of the long-term benefits of *n*-6 PUFA intake for prevention of type 2 diabetes and cardiovascular disease^56-59^. At least with respect to the fat cell membrane lipidome, “you are what you eat”. The diet-dependent composition of adipocyte membranes determines whether mice undergo proper tissue expansion or develop dysfunctional fat storage and metabolic disease in response to chronic overnutrition.

Multiple lines of evidence suggest a pivotal role for arachidonoyl PE in LPCAT3-dependent regulation of LD size. First, the relative abundance of arachidonate present in the PE lipid class far exceeds that in the PC class (∼10:1). Second, the LPCAT3 inhibitor (*R*)-HTS-3 potently suppressed arachidonoyl PE levels and TG storage in cultured adipocytes, with minor effect on arachidonoyl PC. Third, LD-associated membranes isolated from adipocytes displayed a specific enrichment of arachidonoyl PE relative to the ER, in a manner dependent on LPCAT3. Fourth, arachidonoyl PE partitioned to ER–LD junctions in a computational model designed to mimic nascent LD assembly. In line with our shotgun lipidomic analyses, arachidonoyl PC was evenly distributed across the ER bilayer and budding lens. In mammals, PC is synthesized either through the Kennedy (CDP-choline) pathway or via PEMT-dependent methylation of PE to PC^60^. Notably, earlier work has shown that PEMT-derived PC preferentially contains PUFA chains in the *sn*-2 position^61^. While PEMT is mainly expressed in the liver, its activity has also been detected in adipose tissue^62^. Hence, it has yet to be established whether arachidonoyl PC in WAT is generated directly through the Lands cycle or indirectly via the PEMT pathway.

The role of conically shaped lipids such as diacylglycerol and phosphatidic acid in TG nucleation and lens formation is well-documented^63-65^. A possible role for PE in LD biogenesis has received less attention, despite it being the most ubiquitous non-bilayer lipid in cell membranes. Recent *in silico*-based studies have suggested that the negative intrinsic curvature of PE favours “non-budding” conditions, thereby supporting the formation of large LDs^48,49^. However, these studies did not specifically consider the effect of distinct PE fatty acyl-chain conformations on LD size regulation. We add to existing knowledge by demonstrating that PE specifically partitions to the ER–LD interface and that its acyl-chain profile determines LD budding size. The precise biophysical mechanism(s) by which arachidonoyl PE outperforms oleoyl PE in clustering large TG quantities has yet to be fully elucidated. According to our computational analyses, oleoyl phospholipids manifest strong attractions to the triolein core, probably due to their structurally identical acyl-chain profiles. We predict that such high affinity may simply be incompatible with optimal triolein clustering within the ER bilayer. Alternative, but not mutually exclusive, mechanisms through which highly disordered PUFA chains could promote LD budding size include the lowering of membrane bending rigidity, stabilizing the ER–LD neck and/or differentially altering bilayer/monolayer surface tensions to favour TG clustering prior to egress as nascent LDs^47,66,67^.

Our data support a model wherein unilocular adipocytes continuously bud new droplets that merge with large LDs. In a manner akin to CIDE-mediated LD fusion^68^, we propose that increasing LD budding size in adipocytes promotes efficient TG storage by limiting the surface area exposed to abundant cytosolic lipases. This may be particularly important in obese and insulin-resistant states, which are accompanied by impaired PI3K/Akt signalling and elevated cAMP levels^69^. In light of this, the enhanced insulin signalling observed in *Lpcat3*^AKO^ WAT might reflect a compensatory response to limit aberrant lipolysis during times of lipogenic stress. Of interest, prior work from our lab and others has revealed that LPCAT3 can similarly cluster large TG deposits in hepatocytes, which promotes the assembly of VLDL particles^28,29^. Hence, LPCAT3 appears to facilitate TG clustering irrespective of tissue or cell type. The budding directionally might be determined by tissue-specific expression profiles of MTP (liver) and Seipin (adipose). An apparent discrepancy with our study is the observation that *Lpcat3*^KO^ livers display an overt steatotic phenotype^28,29^. However, this can be explained by impaired hepatic VLDL secretion and compensatory budding of cytosolic LDs to mitigate ER stress and lipotoxicity. It is worth noting that ATGL is expressed at much lower levels in the liver, favouring the accrual of LDs regardless of a budding size defect^70,71^.

A fundamental question that arises from the present study is why LPCAT3 activity appears to converge on LD size regulation primarily in unilocular white adipocytes, even though *Lpcat3* is also highly expressed in BAT. One explanation is the existence of cellular mechanisms that enforce a multilocular LD phenotype in brown adipocytes. We have recently discovered a BAT-specific product of the *Clstn3* locus (CLSTN3β) that localizes to ER–LD contact sites and restricts LD size^72^. Optimization of the LD surface area-to-volume ratio in BAT improves the efficiency of TG mobilization in response to cold temperatures. Intriguingly, the *Clstn3* locus is in close spatial proximity to *Lpcat3*, raising the prospect that CLSTN3β has co-evolved with LPCAT3 to limit its propensity to form large LDs in thermogenic fat. Although we have excluded a role for BAT in the *Lpcat3*^AKO^ DIO phenotypes, this does not rule out a cell-intrinsic function of LPCAT3 in brown fat biology. For example, it has been known since the 1970s that *n*-6 PUFAs are selectively enriched in BAT membranes during cold exposure^73^. Further studies will be required to address the physiologic relevance of LPCAT3-dependent membrane remodelling in thermogenesis.

Animal models designed to mimic human lipodystrophies often manifest unexpected ‘lean and healthy’ metabolic phenotypes in response to DIO^18^. Although this is widely believed to be due to a higher metabolic rate, the molecular basis for EE and the tissues coordinating adaptation to dysfunctional fat storage have not been fully explored. Using lipid radiotracer studies and transcriptomic profiling, we have demonstrated that aberrant TG hydrolysis in obese *Lpcat3*^AKO^ iWAT is counterbalanced over time by an adaptive program that engages FA re-esterification, synthesis and oxidation. The onset of TG/FA cycling tracks with increased EE, effectively limiting further body weight gain and lipid spill-over to other tissues. We find that the molecular signature of *Lpcat3*^AKO^ iWAT closely resembles that of lipodystrophic *Cidec-*KO mice^74^. This adaptation to dysfunctional TG storage is therefore likely to be conserved across other models that exhibit defects in LD size and stability. It is also notable that this adaptive WAT-driven EE appears to be independent of classic browning pathways. Recent work has shown that TG/FA cycling has potential for burning excess calories^75^. Future studies into the regulatory mechanisms controlling the rate and/or extent of futile lipid cycling may define targets for therapeutic intervention.

## Acknowledgements

We are grateful to J. Sandhu, C. Priest, and B. Clifford for technical assistance. We also thank other current and former members of the Tontonoz and Tarling-Vallim labs for valuable discussions and sharing reagents. Oxylipin analysis was performed at the RIKEN Center for Integrative Medical Sciences with the assistance of M. Honda. PET/CT was performed at the Crump Institute Preclinical Imaging Technology Center with assistance of S. Xu and M. Tamboline. RNA-seq and sNuc-seq analyses were performed at the Technology Center for Genomics & Bioinformatics. We dedicate this manuscript to the memory of Tom C. Kemper, a beloved friend and colleague. This work was supported by an American Diabetes Association fellowship (1-19-PDF-039 to M.J.T.), grants from the National Institutes of Health (R01DK129276 to P.T.; HL139725 to S.G.Y.), Japan Society for the Promotion of Science abroad, the Osamu Hayaishi Memorial Scholarship for Study Abroad (to Y.S.), the Swiss National Science Foundation (grants 310030_219264 to S.V.), and the European Research Council under European Union’s Horizon 2020 research and innovation program (grant agreement no. 803952, to S.V.). This work was supported by grants of the Swiss National Supercomputing Centre (CSCS) under project ID s1131 and s11876.

## Author Contributions

Conceptualization was done by M.J.T., Y.S., S.V. and P.T. Methodology was developed by M.J.T., Y.S., A.H.B., J.S., L.C., M.C.G., P.H., K.J.W., M.A., D.A.F. and S.V. Formal analysis was carried out by M.J.T., Y.S., A.H.B., J.S., L.C., K.Q., K.J.W., D.P.P., M.A. and S.V. Computational analysis was performed by A.H.B. and J.S. Lipidomics was carried out by K.J.W., B.S., D.P.P. and D.A.F. Investigation was done by M.J.T., Y.S., A.H.B., J.S., L.C., A.F., K.Q., J.P.K., S.D.L., Y.G., X.X., J.G., J.J.M., T.A.W., K.J.W., D.P.P., and M.A. Resources were provided by M.C.G., P.H., C.P., A.J.L., K.J.W., B.S., A.R., N.M., M.A., B.F.C., S.G.Y., D.A.F., R.Z., S.V. and P.T. Data were curated by M.J.T., Y.S., A.H.B., J.S, T.A.W., K.J.W., M.A., S.V. and P.T. The original draft was written by M.J.T., Y.S. and P.T. Review and editing of the draft were done by M.J.T., Y.S., A.H.B., J.S., L.C., M.C.G., P.H., A.F., K.Q., J.P.K., S.D.L., Y.G., X.X., J.G., J.J.M., T.A.W., C.P., A.J.L. K.J.W., B.S., D.P.P., A.R., N.M., B.F.C., M.A., S.G.Y., D.A.F., R.Z., S.V., and P.T. Supervision was the responsibility of M.J.T., Y.S., S.V., and P.T.

## Extended Data Figure Legends

**Extended Data Fig. 1.**
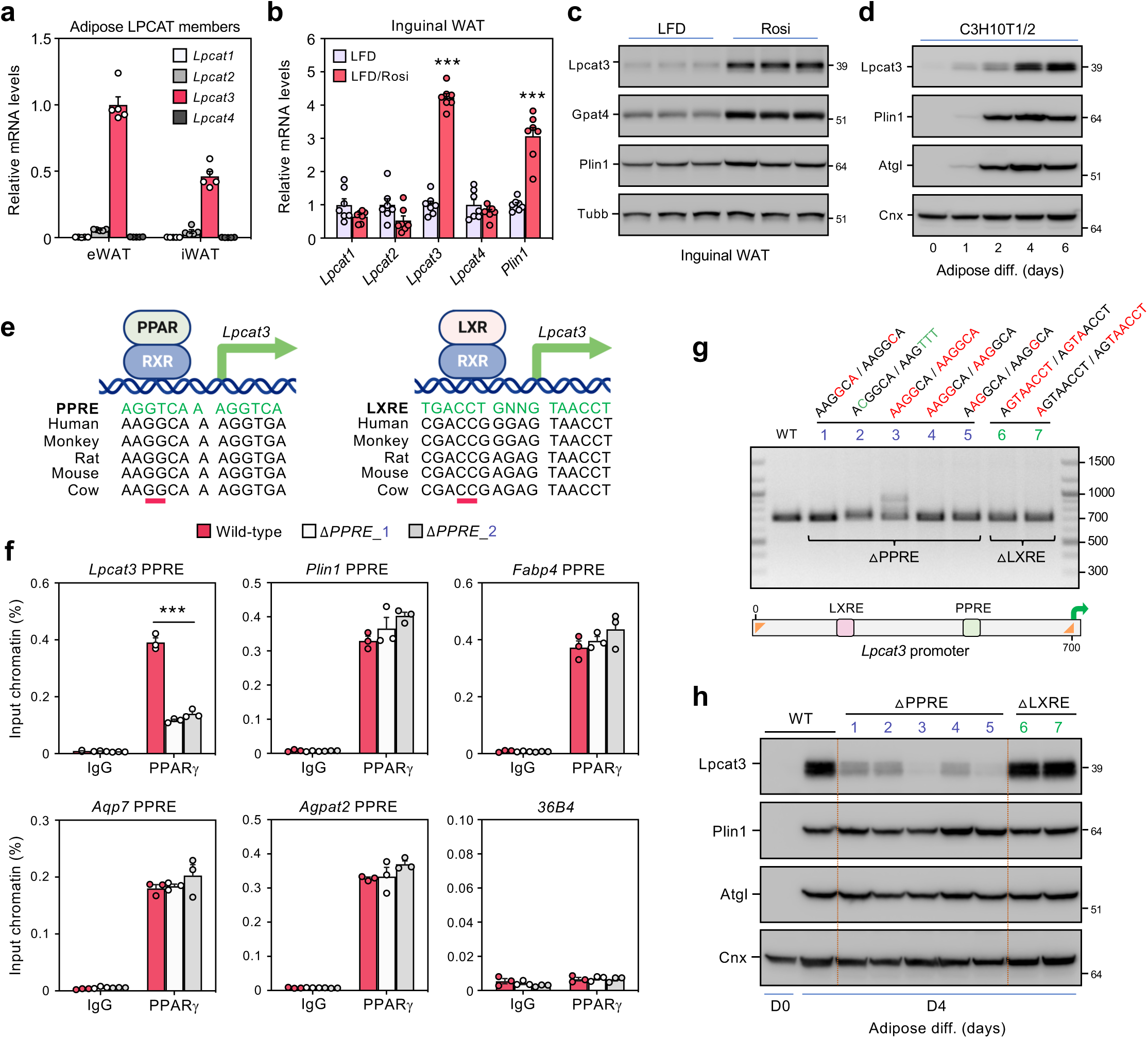
*Lpcat3* is an evolutionarily conserved PPARγ target gene. (**a**) qPCR analysis of *Lpcat1-4* mRNA levels in eWAT and iWAT of 12-week-old chow-fed wild-type mice (*n* = 5, 5), and (**b**) in iWAT from 12-week-old wild-type mice fed low-fat diet (LFD) with or without the specific PPARγ agonist Rosiglitazone (Rosi; 50 mg kg^-1^) for 14 days (*n* = 7, 7). (**c**) Immunoblot analysis of LPCAT3 protein levels in iWAT lysates from (b). (**d**) Immunoblot analysis of LPCAT3 protein levels and other adipogenic markers during the course of 10T1/2 adipocyte differentiation. (**e**) Sequence alignment of conserved PPAR and LXR response elements in the Lpcat3 promoter across multiple species. The PAM sequence of sgRNAs targeting *Lpcat3*-PPRE and LXRE are underlined in red. (**f**) ChIP-qPCR analysis of PPARγ binding to *Lpcat3*, *Plin1*, *Fabp4*, *Agpat2*, or *Aqp7* elements in WT and **△**PPRE 10T1/2 adipocytes (*n* = 3). *36B4* served as a negative ctrl (**g**) PCR amplification of a 700-bp genomic segment flanking PPRE/LXRE sites in the *Lpcat3* promoter region of wild-type, **△**PPRE, or **△**LXRE 10T1/2 clonal cell lines, showing no major deletions introduced by CRISPR/Cas9 gene editing. Oligonucleotide anneal sites in *Lpcat3* promoter are shown in orange, indels in green (insertions) and red (deletions). (**h**) Immunoblot analysis of Lpcat3 protein levels in WT, **△**PPRE, and **△**LXRE 10T1/2 adipocytes after 4 days of differentiation. Data are presented as mean ± SEM. \*\*\**P* < 0.001 by multiple *t*-tests with Holm-Sidak’s correction (**b**,**f**).

**Extended Data Fig. 2.**
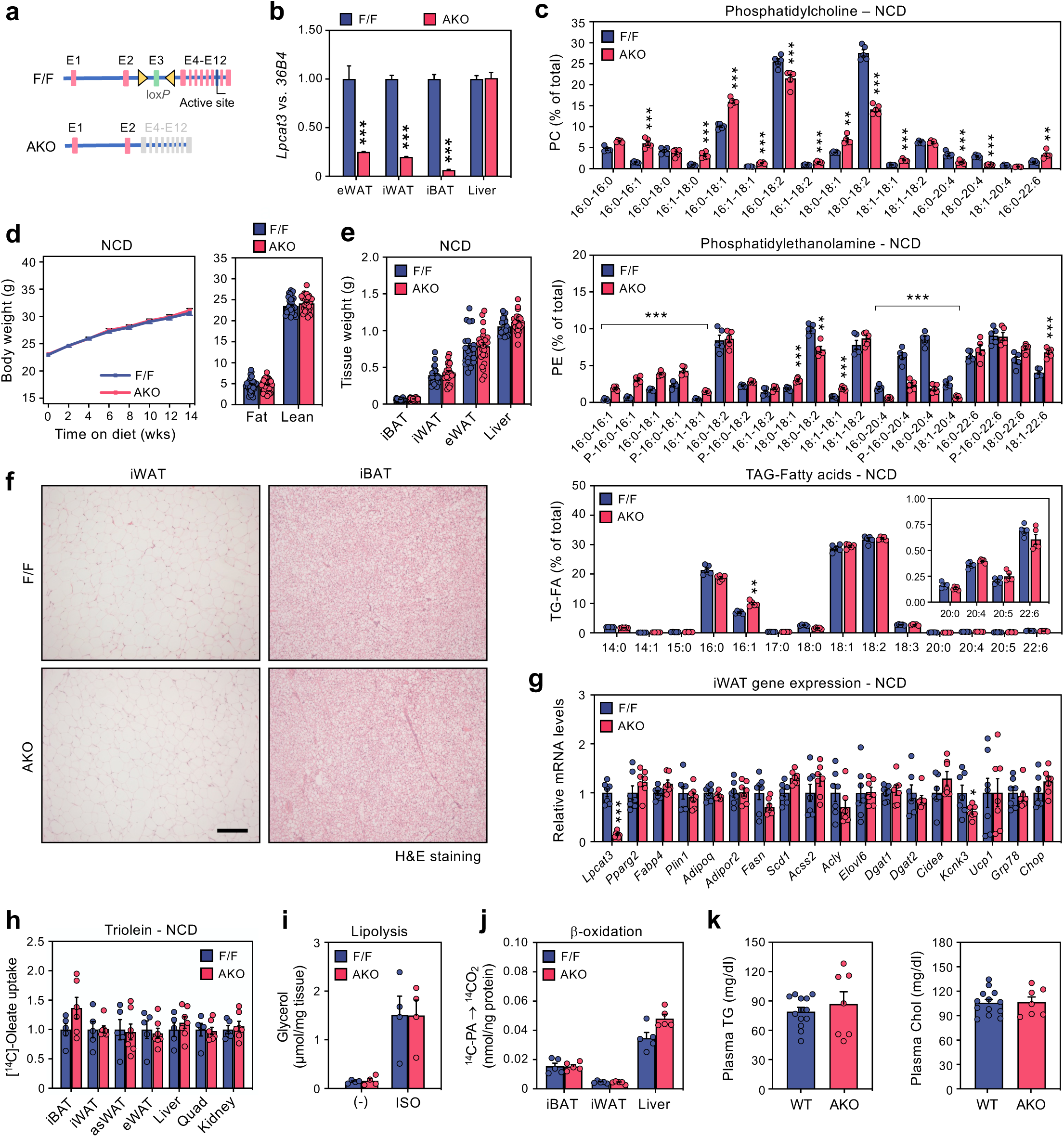
*Lpcat3*^AKO^ mice display no metabolic phenotype under standard laboratory conditions. (**a**) Schematic illustration of *in vivo Lpcat3*^AKO^ strategy. (**b**) qPCR analysis of *Lpcat3* transcript levels in fat depots and liver of 12-week-old NCD-fed control and *Lpcat3*^AKO^ mice (*n* = 3, 3). (**c**) Lipidomic analysis of PC, PE, or TG-FA subspecies in iWAT of NCD-fed *Lpcat3*^AKO^ mice and controls (*n* = 5, 5). (**d**) Body weight (BW) curves and composition of NCD-fed *Lpcat3*^AKO^ mice and controls (*n* = 34, 30). (**e**) Wet weights of distinct fat depots and livers in 22-week-old NCD-fed *Lpcat3*^AKO^ mice and controls (*n* = 23, 25). (**f**) Representative H&E stainings of iBAT and iWAT (scale bar, 100 µm). (**G**) qPCR analysis of iWAT of NCD-fed *Lpcat3*^AKO^ mice and controls (*n* = 7, 7). (**h**) ^14^C-oleate uptake in fat depots and peripheral insulin-target tissues of 18-weeks-old NCD-fed *Lpcat3*^AKO^ mice and controls (*n* = 5, 7). Radioactivity (counts per min, CPM) was calculated in the extracted lipids from whole-organs 4 h post-gavage with ^14^C-Triolein and normalized for g/tissue. (**i**) *Ex vivo* lipolysis in freshly dissected iWAT explants from NCD-fed control and *Lpcat3*^AKO^ mice (*n* = 4, 5). Secreted glycerol was analysed in the supernatant under basal and stimulated (2 *µ*M isoproterenol, ISO) lipolytic conditions. (**j**) *Ex vivo* β-oxidation in crude iWAT, iBAT and liver lysates, as judged by the conversion of ^14^C-palmitic acid (PA) into ^14^CO_2_ (*n* = 5/group). (**k**) Plasma triglycerides and cholesterol in 18-weeks-old NCD-fed *Lpcat3*^AKO^ mice and controls (*n* = 13, 7). Data are presented as mean ± SEM. \*\**P* < 0.01, \*\*\**P* < 0.001 by multiple *t*-tests with Holm-Sidak’s multiple corrections on the CLR-transformed values for each lipid class (**c**), or two-sided Welch’s *t*-test (**b**,**g**).

**Extended Data Fig. 3.**
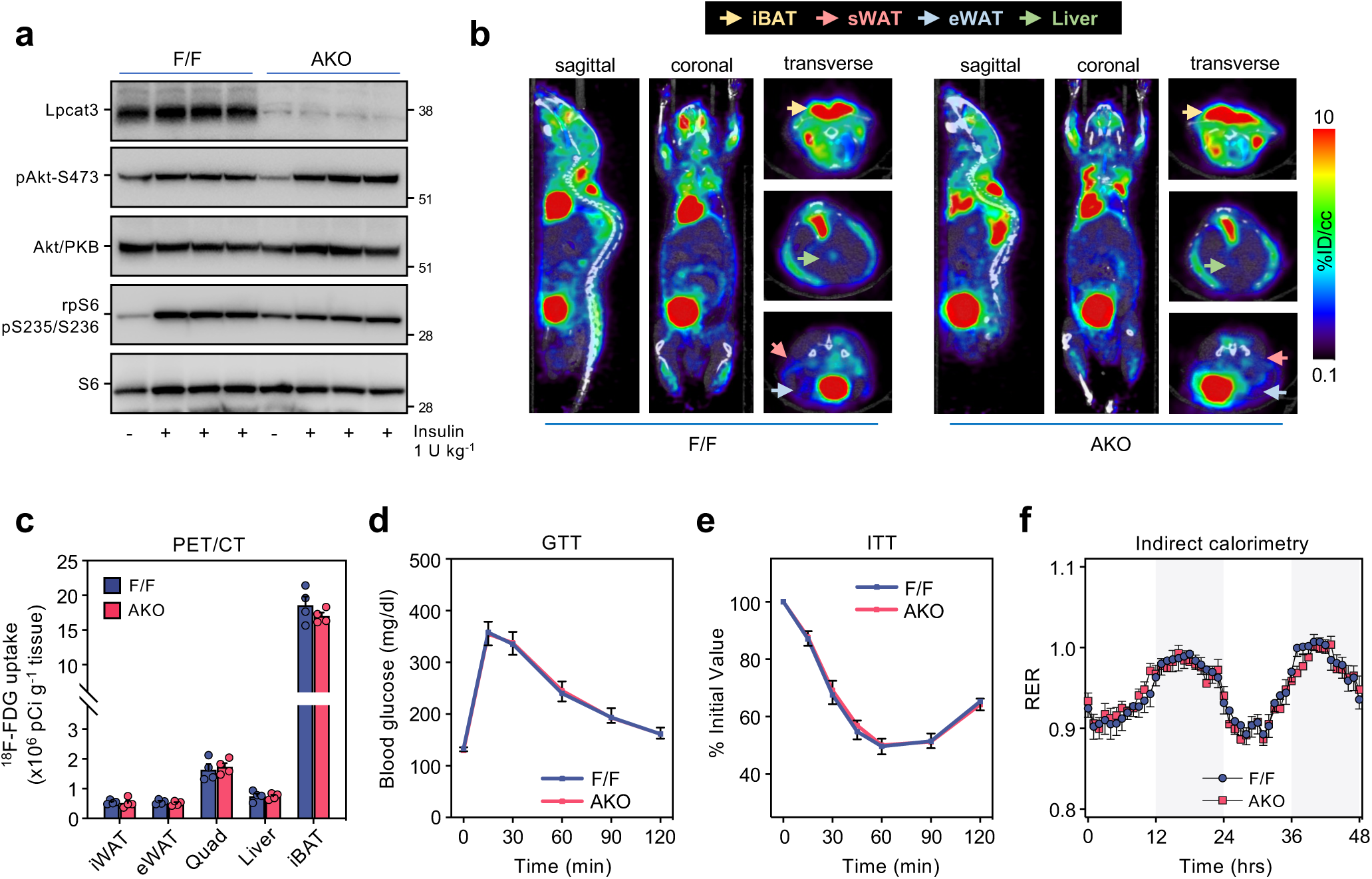
*Lpcat3*^AKO^ mice have no defects in adipocyte and systemic glucose homeostasis. (**a**) Immunoblot analysis of phosphorylated Akt/PKB and rpS6 in iWAT lysates of 12-week-old NCD-fed *Lpcat3*^AKO^ mice and controls after i.p. injection with insulin (1 U kg^-1^). (**b**) Coronal, sagittal, and transverse views of control and *Lpcat3*^AKO^ mice undergoing PET/CT 1 h post-injection with ^18^F-FDG and insulin (1 U kg^-1^). (**c**) ^18^F-FDG uptake (× 10^6^ pCi g^-1^ tissue) in the indicated insulin-target tissues (*n* = 4). (**d**) glucose tolerance tests (1.5 g kg^-1^, i.p.) and (**e**) insulin tolerance tests (0.75 U kg^-1^, i.p.) performed on 18-week-old *Lpcat3*^AKO^ mice and controls (*n* = 9, 10). (**f**) Respiratory exchange ratios (RER) of 18-week-old NCD-fed *Lpcat3*^AKO^ mice and controls (*n* = 16, 15) were monitored over a period of 48 h in Oxymax metabolic cages (12 h light/dark cycles). Data are presented as mean ± SEM.

**Extended Data Fig. 4.**
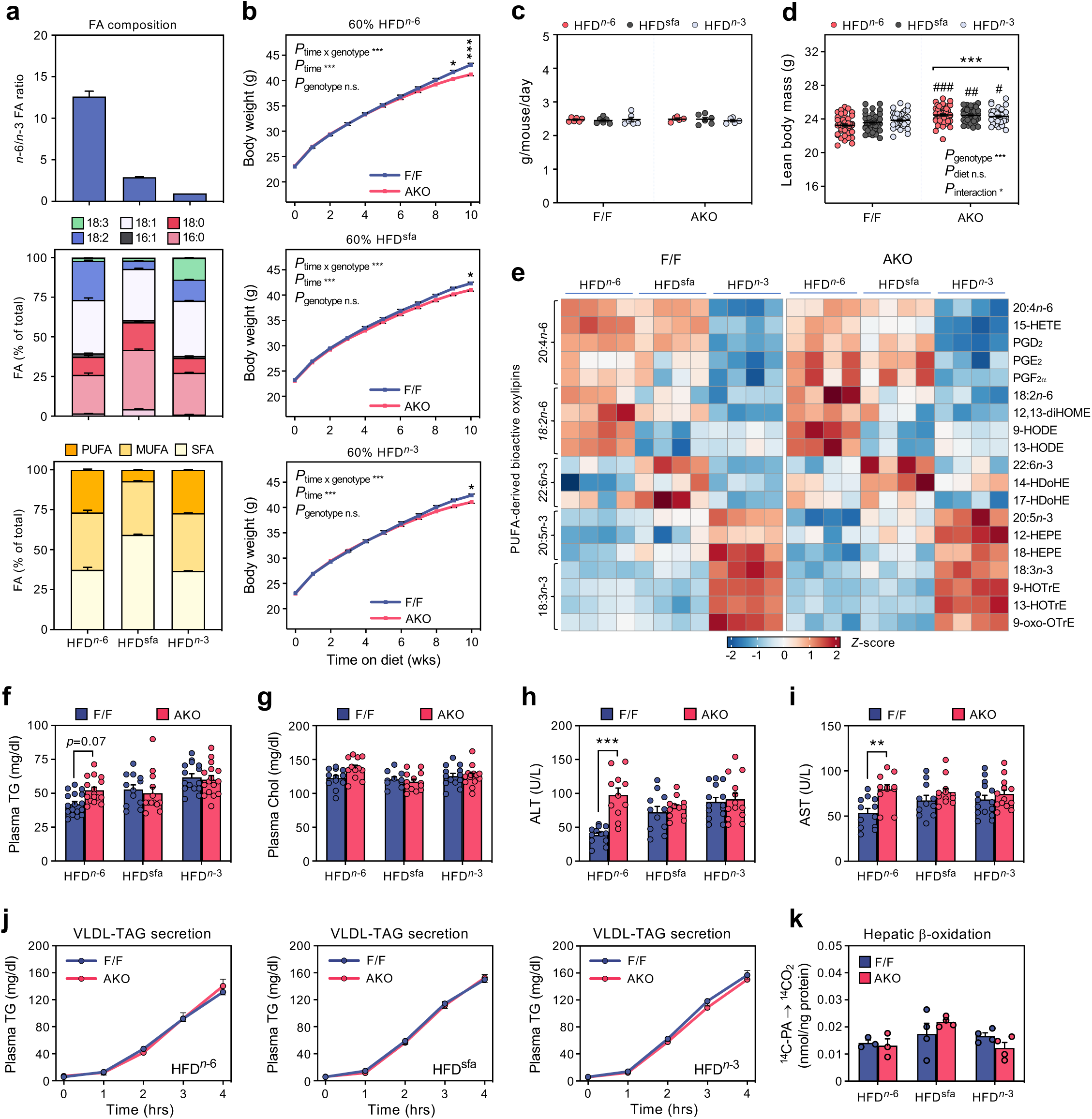
Systemic consequences of lowering adipose membrane *n*-6 PUFA levels in DIO. (**a**) GS-MS analysis of dietary fatty acid profiles and presented as *n*-6/*n*-3 FA ratio, FA species (% of total) or FA class/type (sfa:mufa:pufa). (**b**) Body weight gain curves of 18-week-old *Lpcat3*^AKO^ mice and controls fed HFD*^n^*^-6^ (*n* = 45, 42); HFD^sfa^ (*n* = 46, 38); or HFD^*n*-3^ (*n* = 46, 34) for 10 weeks. (**c**) Average daily food intake of singly-housed *Lpcat3*^AKO^ mice and controls fed the indicated HFDs, recorded over the last 5 weeks of HFD-feeding (*n* = 5–6/group). (**d**) EchoMRI analysis of lean mass in *Lpcat3*^AKO^ mice and controls fed the indicated HFDs for 10 weeks. (**e**) Oxylipins were measured in the inguinal adipocyte fraction isolated from *Lpcat3*^AKO^ mice and controls fed the indicated HFDs for 10 weeks (*n* = 4/group). Oxylipin abundances were transformed using a centered-log ratio (CLR) on the closed composition for each sample. CLR-transformed abundances for each lipid were normalized to allow for sample-wise comparisons. (**f-i**) Plasma (**f**) TGs, (**g**) cholesterol, (**h**) alanine aminotransferase (ALT), and (**i**) aspartate transaminase (AST) levels in *Lpcat3*^AKO^ mice and controls fed the indicated HFDs for 10 weeks (*n* = 10–17/group). (**j**) Very-low density lipoprotein (VLDL) secretion in 14-week-old *Lpcat3*^AKO^ mice and controls fed the indicated HFDs for 6 weeks (*n* = 5–6/group). Mice were fasted for 12 h prior to i.p. injection with the LPL inhibitor poloxamer-407 (1 g kg^-1^). Plasma TGs were measured in blood collected retro-orbitally at the indicated times post-injection. (**k**) β-oxidation in liver lysates, as judged by the conversion of ^14^C-palmitic acid (PA) into ^14^CO_2_ (*n* = 3–4/group). Data are presented as mean ± SEM. \**P* < 0.05, \*\*\**P* < 0.001 using a two-way RM ANOVA with Sidak’s multiple comparisons test (**b**). \*\*\**P* < 0.001 vs. HFD*^n^*^-6^-fed control; ^#^*P* < 0.05, ^##^*P* < 0.01, ^###^*P* < 0.001 vs. HFD^sfa^; same genotype, a two-way RM ANOVA followed by Tukey’s multiple comparisons test (**d**), \*\**P* < 0.01, \*\*\**P* < 0.001 by multiple *t*-tests with Holm-Sidak’s correction (**h**,**i**).

**Extended Data Fig. 5.**
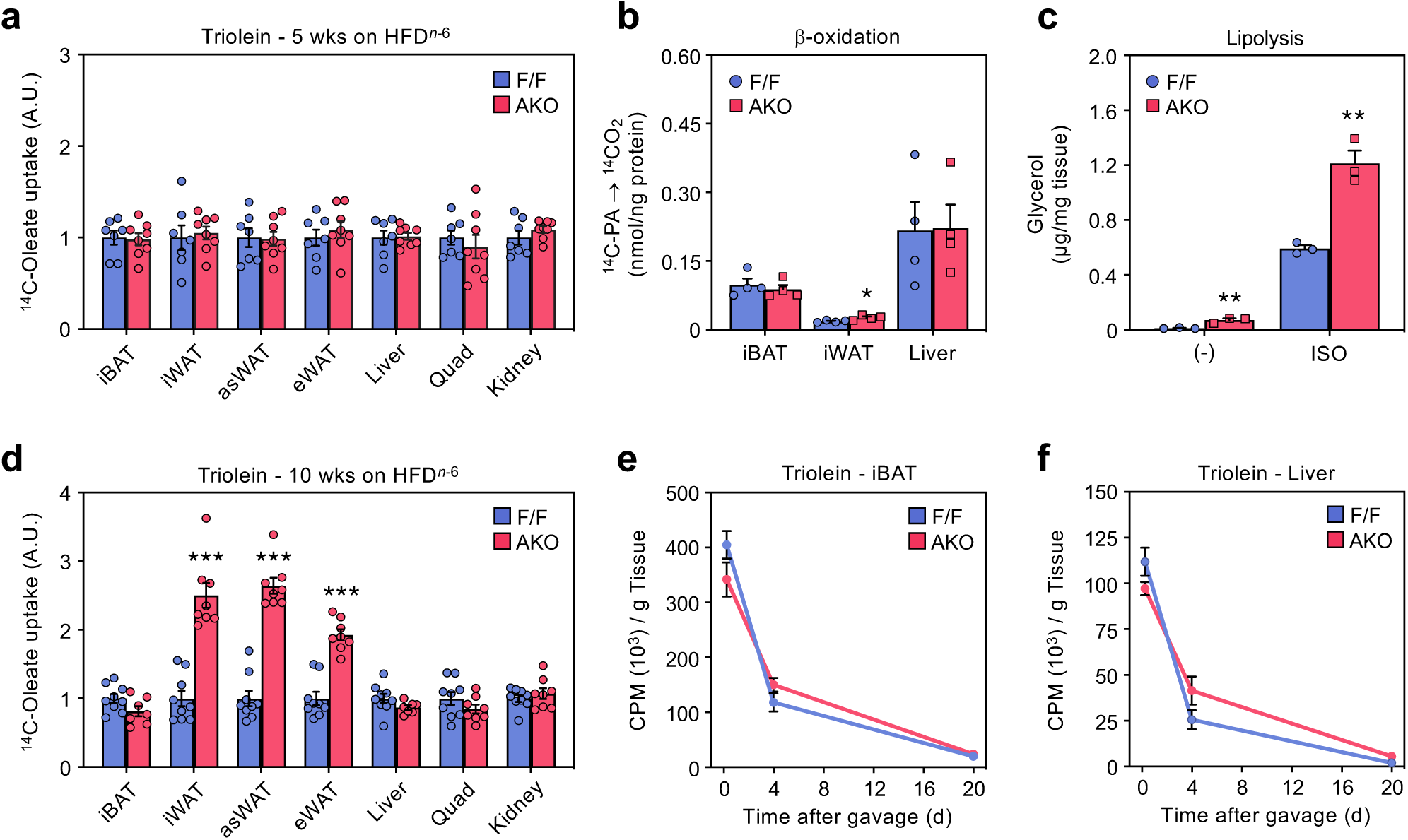
Lipolytic dysregulation in obese *Lpcat3*^AKO^ mice. (**a**)^14^C-oleate uptake in distinct fat depots and peripheral insulin-target tissues from control and *Lpcat3*^AKO^ mice fed a HFD*^n^*^-6^ for 5 weeks (*n* = 7, 8). Radioactivity (counts per min, CPM) was calculated in extracted lipids from whole-organs 4 h post-gavage with ^14^C-Triolein and normalized for g/tissue. (**b**) *Ex vivo* β-oxidation rates in crude iWAT, iBAT, or liver lysates from *Lpcat3*^AKO^ mice and controls fed HFD*^n^*^-6^ for 5 weeks, as assessed by the conversion of ^14^C-PA into ^14^CO_2_ (*n* = 4, 4). (**c**) *Ex vivo* lipolysis in iWAT explants freshly dissected from *Lpcat3*^AKO^ mice and controls fed HFD*^n^*^-6^ for 10 weeks (*n* = 3, 3). Released glycerol was determined under basal or stimulated (2 *µ*M ISO) lipolytic states. (**d**) ^14^C-oleate uptake in the indicated tissues from *Lpcat3*^AKO^ mice and controls fed a HFD*^n^*^-6^ for 10 weeks (*n* = 8, 9). (**e**,**f**) ^14^C-oleate uptake and turnover in (**e**) iBAT and (**f**) the liver from control and *Lpcat3*^AKO^ mice fed HFD*^n^*^-6^ for 10 weeks (*n* = 6–9/group). Radioactivity (counts per min, CPM) was calculated in the extracted lipids from whole-organs 4 h, 4 d, or 20 d post-gavage with ^14^C-Triolein and normalized for g/tissue. Data are presented as mean ± SEM. \*\**P* < 0.01, \*\*\**P* < 0.001 by multiple *t*-tests with Holm-Sidak’s correction (**c**-**d**).

**Extended Data Fig. 6.**
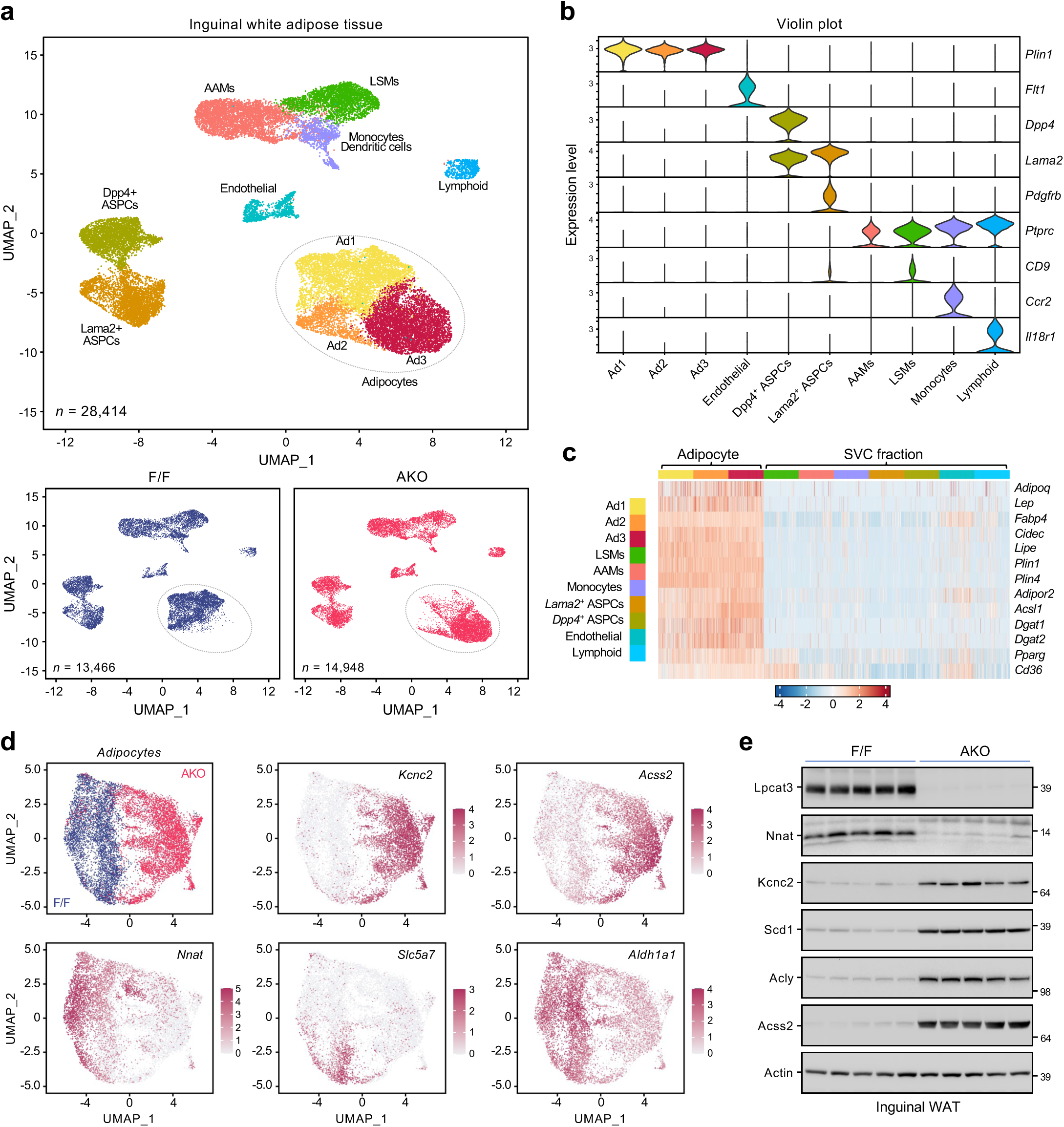
Membrane *n*-6 PUFA levels inversely correlate with adipocyte DNL in obesity. (**a**) UMAP projection of 28,414 sequenced iWAT nuclei or split by genotype (13,466 and 14,948 nuclei for 18-week-old control *vs*. *Lpcat3*^AKO^ mice fed a HFD*^n^*^-6^ for 10 weeks. (**b**) Violin plots (clusters as columns, genes as rows) of cluster-specific markers. (**c**) Heatmap of normalized expression values of pan-adipogenic markers in inguinal adipocytes and the stromal vascular cell fraction. Alternatively activated macrophages (AAMs), lipid-scavenging macrophages (LSMs), adipose stem and progenitor cells (ASPCs). (**d**) *t-*SNE-plots of Ad subtype and genotype-specific markers. (**e**) Immunoblot analysis of DNL markers in iWAT of control and *Lpcat3*^AKO^ mice fed HFD*^n^*^-6^ for 10 weeks (*n* = 5, 5).

**Extended Data Fig. 7.**
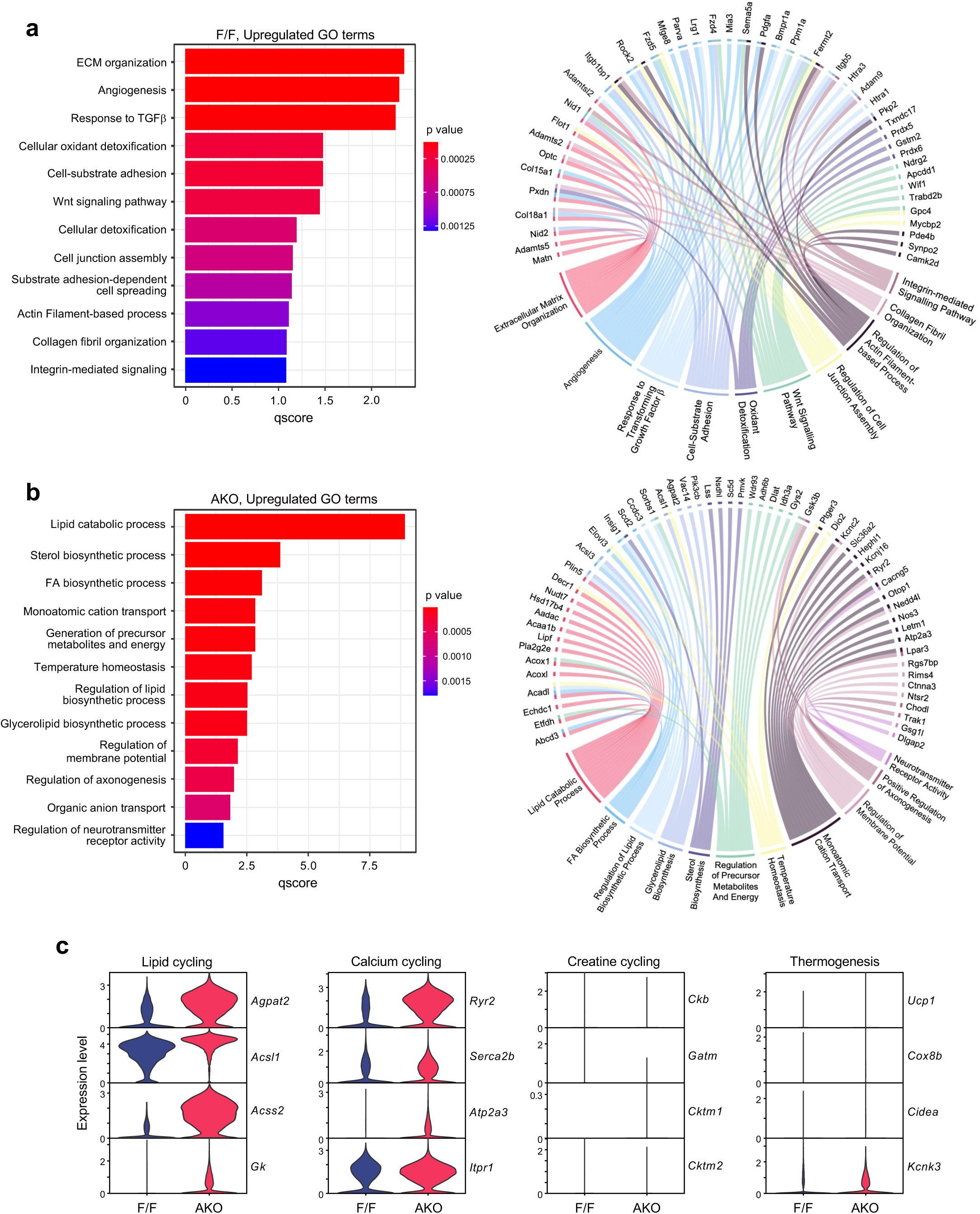
The adaptive *Lpcat3*^AKO^ iWAT signature in DIO is enriched for lipid cycling. (**a**,**b**) Gene ontology (GO) and circos plot of upregulated genes in iWAT of (**a**) control and (**b**) *Lpcat3*^AKO^ mice fed a HFD*^n^*^-6^ for 10 weeks. GO terms were based on the top *Nnat/Aldh1a1* and *Kcnc2* co-expressing iWAT transcripts in integrated RNA-seq and sNuc-seq datasets. (**c**) Violin plots of representative markers for UCP1-dependent and -independent futile cycles.

**Extended Data Fig. 8.**
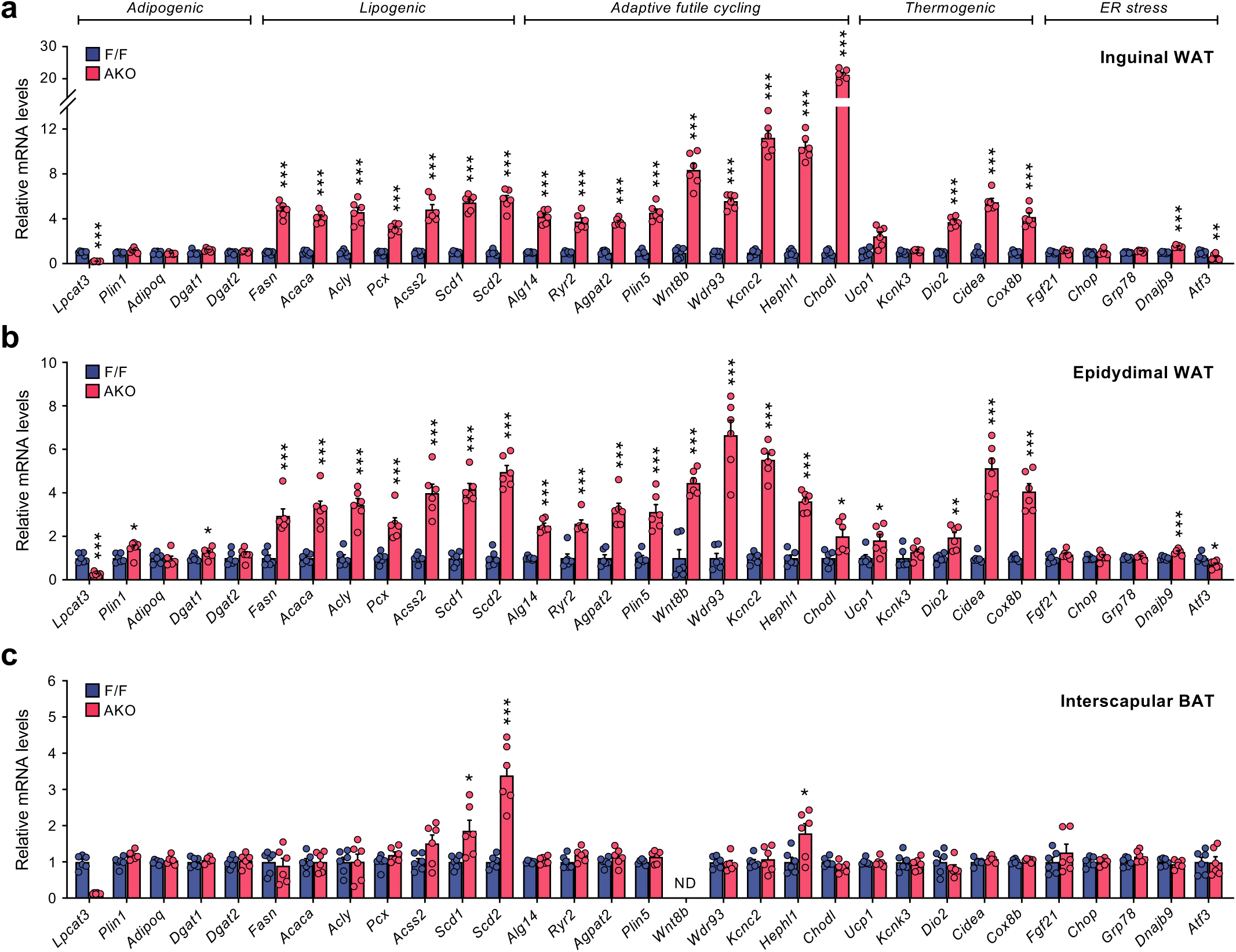
The adaptive *Lpcat3*^AKO^ response to DIO is WAT-selective. (**a**-**c**) qPCR analysis of pan-adipogenic, DNL, lipid cycling, thermogenic, and ER stress-related gene programs in (**a**) iWAT, (**b**) eWAT, and (**c**) iBAT depots of 18-week-old *Lpcat3*^AKO^ mice and controls fed a HFD*^n^*^-6^ for 10 weeks (*n* = 6, 6). Data are presented as mean ± SEM. \**P* < 0.05, \*\**P* < 0.01, \*\*\**P* < 0.001 by multiple *t*-tests with Holm-Sidak correction (**a**-**c**).

**Extended Data Fig. 9.**
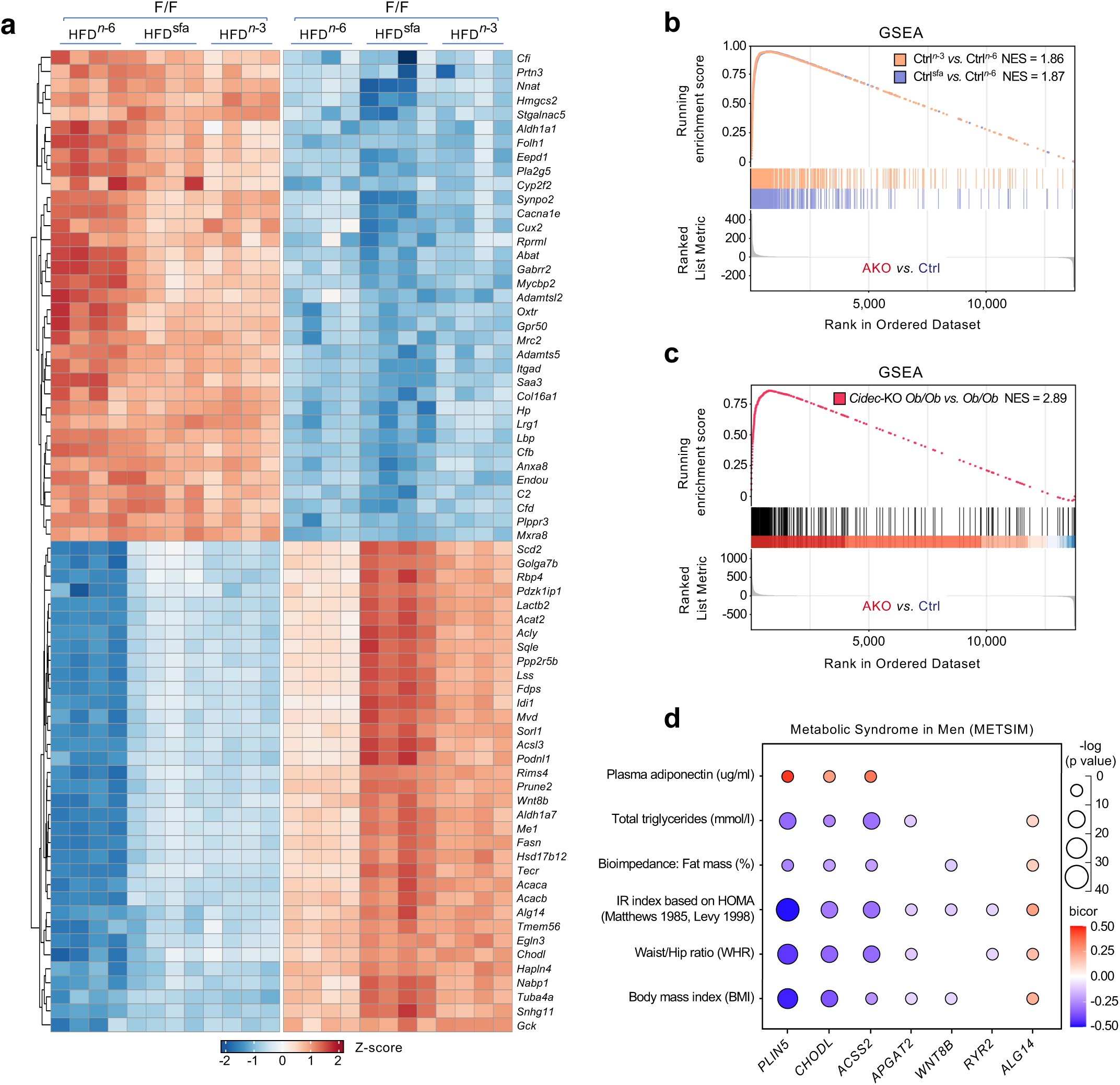
Dietary *n*-6 PUFA restriction or *Cidec*^KO^ mirror *Lpcat3*^AKO^ adaptation to DIO. (**a**) Heatmap representation of the top 70 differentially expressed genes (35 up or down) in iWAT depots of 18-week-old *Lpcat3*^AKO^ mice and controls fed the indicated HFDs for 10 weeks. (**b**,**c**) GSEAs of the top 500 upregulated genes in (**b**) iWAT from HFD^*n*-3^/HFD^sfa^-fed controls *vs*. HFD^n-6^-fed controls or (**c**) from eWAT of *ob/ob Cidec*-KO mice vs. *ob/ob* controls^74^. The ranked list metric is the product of log_2_ (fold-change)* – log_10_ (adjusted *P* value) from the differentially expressed genes in iWAT from HFD-fed *Lpcat3*^AKO^ mice *vs.* controls (normalized enrichment score, NES). (**d**) Correlation matrix depicting the association between key *Lpcat3*^AKO^ markers and human clinical traits in the METSIM study^76^. Node color and size reflect correlation directionality and *p* value. Correlations were analysed by midweight bicorrelation coefficient or corrected *p* value using the *R* package WGCNA.

**Extended Data Fig. 10.**
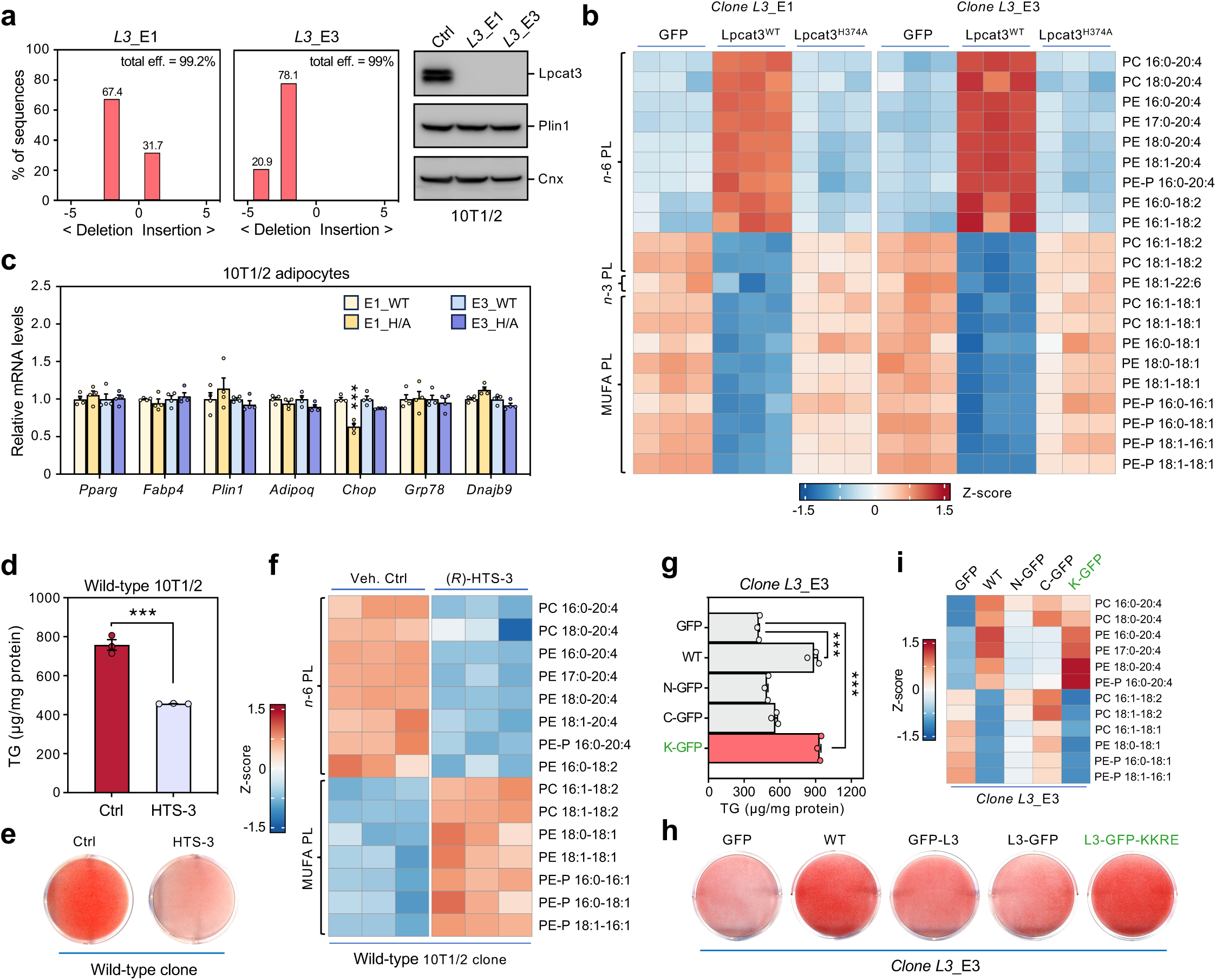
Characterization of *Lpcat3*-null 10T1/2 adipocytes. (**a**) TIDE analysis (left) and immunoblot analysis of LPCAT3 protein levels (right) verifying *Lpcat3*-KO in 10T1/2 adipocyte clonal cell lines transfected with single-guide RNAs targeting *Lpcat3* exon 1 (*L3*_E1) or exon 3 (*L3*_E3). (**b**) Shotgun-lipidomic analysis and hierarchical clustering of PC/PE species in *Lpcat3*-null adipocytes stably expressing GFP, Lpcat3^WT^, or Lpcat3^H374A^ (day 7 of differentiation). (**c**) qPCR analysis of pan-adipogenic markers in *Lpcat3*-KO adipocytes expressing Lpcat3^WT^ or Lpcat3^H374A^ and differentiated for 6 days (*n* = 4, 4). (**d**) TG content (*n* = 3, 3), (**e**) ORO staining, and (**f**) lipidomic analysis of a wild-type 10T1/2 clone differentiated in the presence of 0.1% DMSO (vehicle, veh ctrl) or LPCAT3 inhibitor (*R*)-HTS-3 (10 *µ*M) for 6 days. (**g**) TG content (*n* = 3, 3), (**h**) ORO staining, and (**i**) lipidomic analysis of *Lpcat3*-null adipocytes stably expressing either GFP control or the indicated GFP-tagged Lpcat3 fusion sequences (day 7 of differentiation). Data are presented as mean ± SEM. \*\*\**P* < 0.001 by multiple *t*-tests with Holm-Sidak’s correction (**c**) or Welch’s t-test (**d**,**g**).

**Extended Data Fig. 11.**
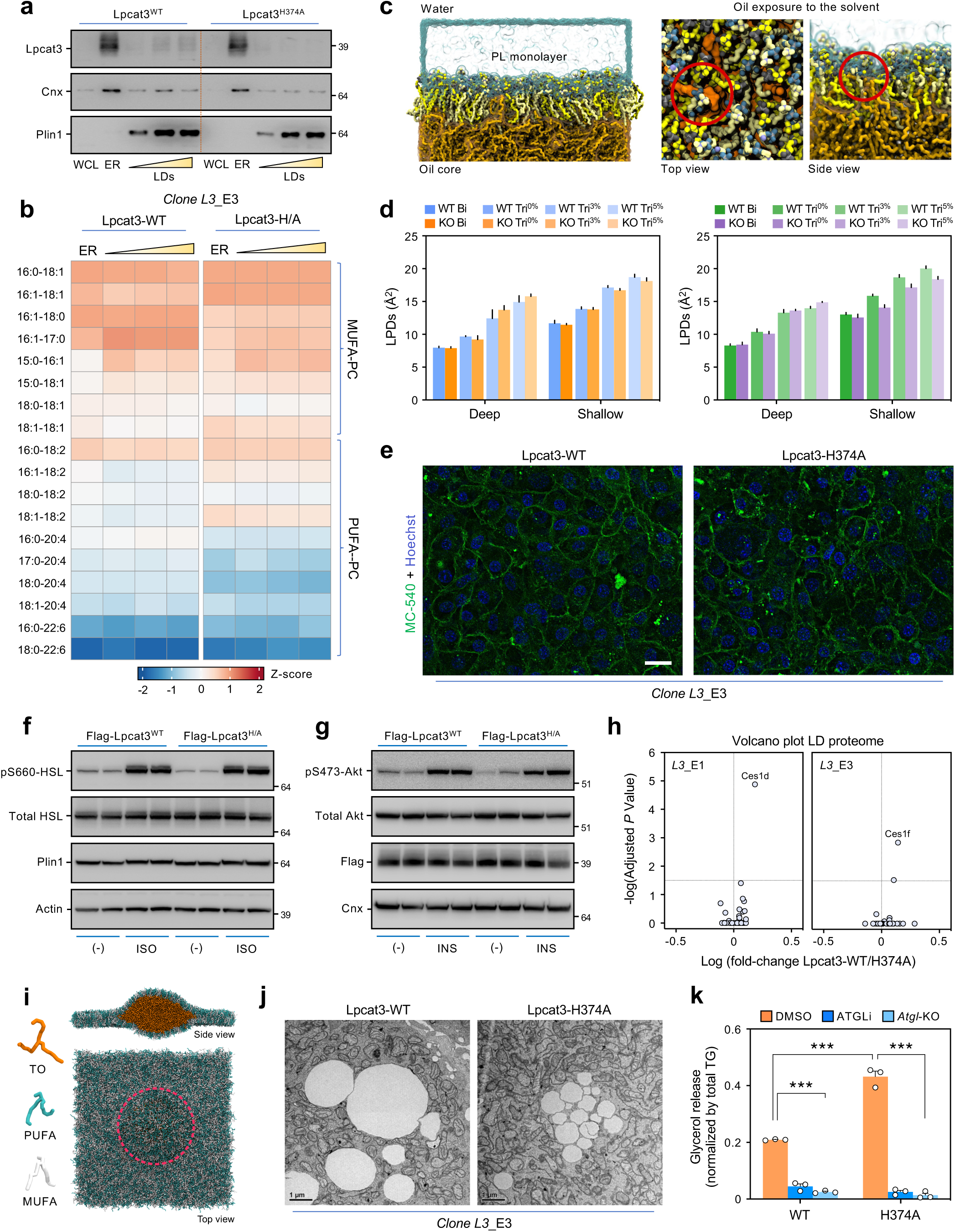
Adipocyte Lpcat3 activity does not affect gross membrane lipid-packing. (**a**) Immunoblot analysis of purified ER membranes and buoyant LD/ER-enriched fractions with small/nascent (20k × *g*), medium (8k × *g*), or large (500 × *g*) LDs from *Lpcat3*-null adipocytes stably expressing Lpcat3^WT^ or Lpcat3^H374A^ (day 6 of differentiation). Similar amounts of protein from each fraction were loaded onto gel (as estimated by silver stain). (**b**) Shotgun-lipidomic analysis of PC composition in ER or LD/ER-enriched fractions from *Lpcat3*-null adipocytes expressing Lpcat3^WT^ or Lpcat3^H374A^ (day 6 of differentiation). Data are presented as the distribution of PC molecular species (% of total PC) within each sample. (**c**) Schematic depiction of a trilayer system containing a bulk oil phase flanked by a phospholipid monolayer and solvated with water (left), and top/side views of trilayers exhibiting LPDs that expose the core to the solvent (right). (**d**) Quantification of deep/shallow LPDs in model bilayers and LD-like trilayers (± 3 or 5% surface tension, ST). WT and *Lpcat3*-KO compositions are either complex (left) or simplified (right) representations of the lipidomic analysis (Supplementary Table 5). (**e**) Live-cell confocal imaging of lipid-packing in *Lpcat3*-null adipocytes expressing Lpcat3^WT^ or Lpcat3^H374A^ (scale bar, 20 µm). Adipocytes were stained with Hoechst and the phospholipid packing probe MC-540 (300 nM) 15 min prior to imaging. (**f**,**g**) Immunoblot analysis of *Lpcat3*-null adipocytes expressing Lpcat3^WT^ or Lpcat3^H374A^ and stimulated with (**f**) ± isoproterenol (ISO, 10 *µ*M), or (**g**) ± insulin (100 nM) for 15 min. (**h**) Tandem mass tag (TMT) labelling performed on buoyant 8,000 × *g* LD-enriched fractions isolated from *Lpcat3*-KO adipocytes expressing Lpcat3^WT^ or Lpcat3^H374A^. The TMT data were overlaid with the Uniprot mouse proteome with annotated LD localization. (**i**) Top/side views of a representative snapshot from coarse-grain MD simulations of an ER–LD neck model system. The bilayer/monolayers consist of 50% PUFA (cyan) and 50% MUFA (white) phospholipids, with an embedded triolein (TO; orange) core. (**j**) TEM analysis of *Lpcat3*-null adipocytes expressing Lpcat3^WT^ or Lpcat3^H374A^ (day 3 of differentiation). (**k**) Stimulated lipolysis in *Lpcat3*^KO^ and *Lpcat3*/*Atgl*^DKO^ adipocytes expressing Lpcat3^WT^ or Lpcat3^H374A^ (day 6 of differentiation). Released glycerol was measured following isoproterenol (ISO, 10 *µ*M) stimulation for 2 h and normalized for intracellular TG stores (*n* = 3/group). Where indicated, cells were incubated in the presence of 0.1% DMSO (veh ctrl) or ATGLi (40 *µ*M) for 6 days (*n* = 3/group). Data are presented as mean ± SEM. Protein abundance ratios were compared across groups using Student’s t-test (*P* < 0.01) while controlling for false-discovery rate with Benjamini-Hochberg procedure (**h**), and \*\**P* < 0.001 by multiple *t*-tests with Holm-Sidak’s correction (**k**).

